# Mangrove leaf and root traits and their relation to urbanization

**DOI:** 10.1101/506196

**Authors:** Benjamin Branoff

## Abstract

Root and leaf traits are one means of understanding plant ecophysiological responses to environmental variation and disturbance. In mangroves, both chemical and morphological variations have been recorded in response to changes in inundation, salinity, and nutrient levels. Some have also been noted in urban environments, primarily in response to elevated nutrients and toxic substances. Yet these studies have not attempted to isolate the urban from the non-urban influences on both morphological and chemical traits. This study measured mangrove leaf and root chemical and morphological traits in herbarium samples and in field collected leaves and roots along a quantified urban gradient in three watersheds of Puerto Rico. It then correlated these traits with predictors of surrounding land cover, as well as with metrics of flooding and water chemistry. There were significant lines of evidence leading to an influence of urban sewage and roads on leaf and root traits. Leaf percent nitrogen increased with urbanization and with surface water nitrogen and phosphorus concentrations, but its isotopic content decreased with increasing phosphorus, leading to the hypothesis that both nitrogen and phosphorus are fueling an otherwise co-limited community of mangroves and nitrogen fixing microbes. The most urban site harbored some of the highest metal concentrations, and there was evidence that elevated concentrations primarily influence fine roots. Other morphological traits are more likely explained by both water chemistry and leaf chemistry and should be considered when interpreting the influence of urban landscapes on mangroves. Percent nitrogen in herbarium samples increased at the least urban site, but remained unchanged at the most urban site, reflecting the relative levels of urbanization at the time of the first samples and their subsequent changes. Most metals from herbarium samples decreased or remained unchanged, suggesting regulation and infrastructure have helped to reduce the release of trace metals to the estuaries. Understanding the influence of urbanization in the context of water chemistry and flooding dynamics will aid in the management of these systems as global urbanization continues.

## Introduction

Plant leaf and root morphological and chemical traits are important cues to ecosystem function and environmental conditions (Peter B Reich et al. 1999; Diaz et al. 2004; Westoby and Wright 2006; Bardgett et al. 2014). In leaves, metrics of mass, area, and their ratios of specific leaf area (SLA) and leaf mass per area (LMA) are strong predictors of species-specific nutritional strategies, environmental physio-chemical conditions, and ecosystem community and productivity dynamics (Diaz et al. 2004; Westoby and Wright 2006). Root traits are considerably less understood than leaves, but can still be used to predict variations in plant strategies, environmental conditions, and productivity and community dynamics (Bardgett et al. 2014).

Additional chemical traits, especially of trace metals and stable isotopes, provide further evidence for the physiological mechanisms associated with observed morphological variation in both leaves and roots (Barceló and Poschenrieder 1990; P B Reich et al. 1998; Hultine and Marshall 2000; Bini et al. 2012; Domínguez et al. 2012). Consequently, recent research has begun to use plant traits to understand the ecological responses to spatial variations in chronic disturbances such as those that are often induced by urbanization (Carreras et al. 1996; Verma and Singh 2006; Balasooriya et al. 2009). Other studies have used herbarium samples to show how these traits vary with the temporal progression of urbanization (Dolan et al. 2011; Huang et al. 2016; Palma et al. 2017). Thus, in addition to tracking the present effects of global change on plant ecology, these trait metrics will be a useful tool in predicting the potential changes to ecosystem function in the more urban future (Williams et al. 2009).

Leaf area and SLA, as well stomatal pore surface, conductance, and density have repeatedly been shown to vary with urbanization (Carreras et al. 1996; Hultine and Marshall 2000; Lima et al. 2000; Moraes et al. 2003; Verma and Singh 2006; Balasooriya et al. 2009). This is largely thought to be a toxic or compensatory response to common urban pollutants, such as ozone or sulfuric and nitric oxides. Such contamination is also evident in foliar carbon isotope measurements of δ^13^C, which increase as plants utilize fossil-fuel sourced carbon from urban exhausts (Lichtfouse et al. 2003), as well as in trace metals from sewage or roads (Tomašević et al. 2005; Singh and Agrawal 2007; De Nicola et al. 2008). Root biomass has also been shown to decrease with urbanization, with implications for soil biogeochemistry and nitrogen processing (Gift et al. 2010). This is likely due to higher retention of anthropogenically sourced nitrogen in urban soils (Raciti et al. 2008), which is evident in lower carbon to nitrogen ratios (C:N) and higher δ^15^N in plant tissues (Kendall et al. 2007), and which induces lower belowground biomass allocation (Wilson 1988). In addition to spatial variation along urban gradients, these δ^13^C and δ^15^N isotopic and trace metal measurements in herbarium samples have also been used to reflect changes in urbanization with time (Penuelas and Filella 2002; Martín et al. 2015; Huang et al. 2016). Thus, measuring these traits can provide important information not only on the environmental conditions in urban ecosystems, but also on the plant physiological responses to these conditions and the potential effects at the system level across space and time.

Mangroves, however, are unique from other terrestrial ecosystems in that they inhabit the dynamic physio-chemical environments of coasts. These forested wetlands are subjected to waters of varying temperature, salinity, and flood frequency, which can have marked influence on plant physiology and the resulting leaf and root traits (Cintrón et al. 1978; Ernesto Medina 1999). Both δ^15^N and δ^13^C in mangrove leaves have been shown to respond to urbanization as predicted from studies on terrestrial systems (Branoff 2017a; Branoff 2018a). But these same metrics are also known to change in response to natural variations in nutrient limitation (McKee et al. 2002), and as plants adjust their water-use efficiency in response to non-anthropogenic influences of nutrient limitation, salinity, and flooding (Lin and Sternberg 1992; E Medina and Francisco 1997; Ernesto Medina 1999; Castañeda-Moya et al. 2011). It is also known that trace metals vary in content within mangrove tissues and they have been repeatedly shown to be associated with anthropogenic disturbance (Defew et al. 2005; Geoff R MacFarlane et al. 2007; Qiu et al. 2011). Some of these metals have further been shown to induce stress in mangrove plants, as reflected in enzymatic activity and plant growth (G R MacFarlane and Burchett 2001;

G R MacFarlane and Burchett 2002; G R MacFarlane 2003). Other changes in mangrove foliar and root morphological measurements have also been shown to vary along environmental stress gradients induced by sub-optimal salinity and dissolved oxygen concentrations (McKee 1996; Takemura et al. 2000; Ye et al. 2003; Lira-Medeiros et al. 2010; Flores-de-Santiago et al. 2012; Basyuni et al. 2018), although this has not yet been studied in the context of urbanization. Thus, its possible that mangrove traits will respond to urbanization in similar fashion to terrestrial plants, but environmental variation in nutrients, salinity, and flooding dynamics must also be accounted for.

The mangroves of Puerto Rico have experienced a complex history of social influences over the past two centuries (Martinuzzi et al. 2009). Forest coverage decreased from the colonial period to its low in the 1930’s, when the island’s economy shifted from agricultural to industrial, and has steadily increased to the present day. Spatially, however, these changes have not been uniform. In San Juan, the island’s largest metropolitan area, and also adjacent to the largest mangrove forest on the island, development, dredging, and canalization have markedly transformed the forests beginning in the early 1500’s (United States Army Corps of Engineers 2015). The Caño Martín Peña, which today harbors the most urban mangroves on the island, has arguably been subjected to the most intense and chronic urban disturbances. Bridge building, dredging, canalization, and development encroachment have all occurred during various phases of the canal’s colonial to modern history. Much of this has been un-regulated, resulting in forests today that receive raw human and animal wastes and have been filled with refuse, all of which supposedly alter the channels flow and biogeochemistry. Population density in the surrounding neighborhoods of the canal has increased exponentially from around 400 people/km^2^ in 1910 to roughly 8000 people/km^2^ today (United States Census Bureau, 1910-2000). This is markedly different from the mangroves of Piñones, which have experienced no known major dredging or canalization projected, and whose surrounding population density remained under 500 people/km^2^ until the 1950’s, and whose current population is around 2000 people/km^2^.

This study compares mangrove leaf morphological and chemical traits, as well as root biomass and chemistry from both live trees and herbarium specimens along well-defined urban gradients in Puerto Rico. The influence of urbanization on these traits is analyzed together with that of environmental measurements of surface water chemistry and flooding metrics to elucidate the relative role of each in explaining the observed variation in traits. Such studies will be increasingly useful in the management of mangroves in the twenty-first century, which has already marked a transition from rural to urban human societies, and which will continue to see the relatively rapid urbanization of tropical, sub-tropical and warm temperate coastal areas.

## Methods

Green leaves and both live and dead roots were collected from twenty one-hectare mangrove forest stands across three watersheds in Puerto Rico (Figure 1 Table 1). Forest compositional and structural metrics at each site are described in detail in (Branoff and Martinuzzi 2018). All forests border a waterbody and extend from twenty to one-hundred meters inland. Three plots were selected from each one-hectare site such that all species within the site were represented in the sampling, and that both minimum and maximum distances from the shore were also sampled. Partial sun leaves were collected from each species in the plot at a height of five meters from the ground. A small branch was cut from the tree and six leaves from the 2^nd^ or 3^rd^ youngest cohorts were randomly selected and placed on ice and in a freezer until further processing. In the laboratory, leaf blades were thawed, separated from the petiole, wiped clean, weighed for fresh weight, and measured for morphology metrics of length, width, and area using photos and the ImageJ software (Reinking 2007). Leaves were then dried in a drying oven at 60°C for 48 hours and weighed again for dry weight.

**Figure 1.**
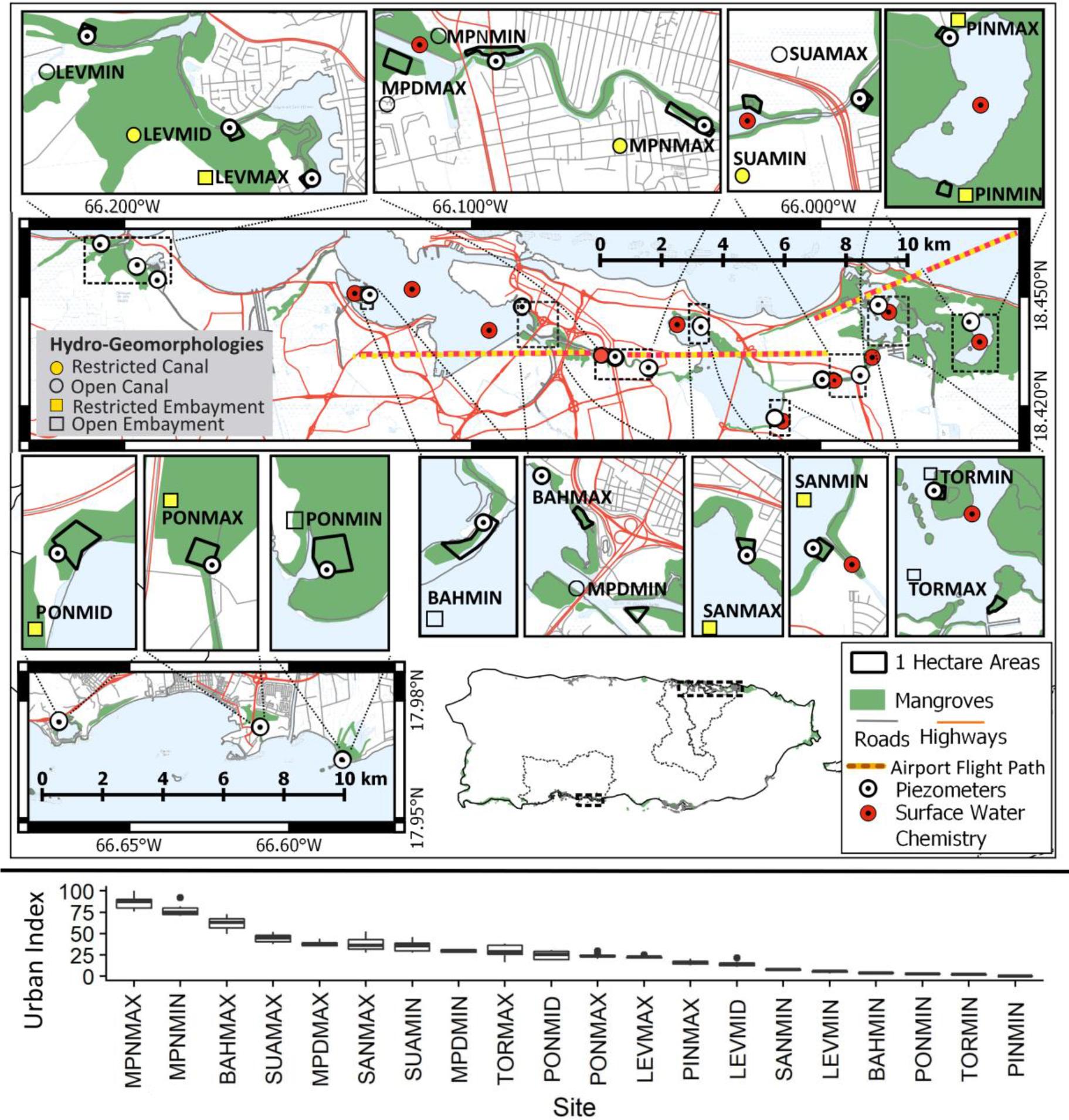
Leaves and roots were taken from twenty one-hectare mangrove forests along a well quantified urban gradient in three watersheds of Puerto Rico. Urbanization was generalized from the urban index, which includes surrounding urban, vegetated, open water, and mangroves land classes, as well as road and population density. An urban index of one-hundred represents the greatest relative level of urbanization. Water-level observations from piezometers were used to construct flooding metrics and surface water chemistry measurements were averaged over five years from a network of stations in the San Juan Bay Estuary. “MAX” and “MIN” postscripts refer to urbanization levels within each water body. BAH is the San Juan Bay, MPD is the dredged portion of the Caño Martín Peña, MPN is the un-dredged portion of the Caño Martín Peña, SAN is the San José lagoon, SUA is the Suárez canal, TOR is the Torrecillas lagoon, PIN is the Piñones lagoon, LEV is Levittown and PON is Ponce.

**Table 1.**
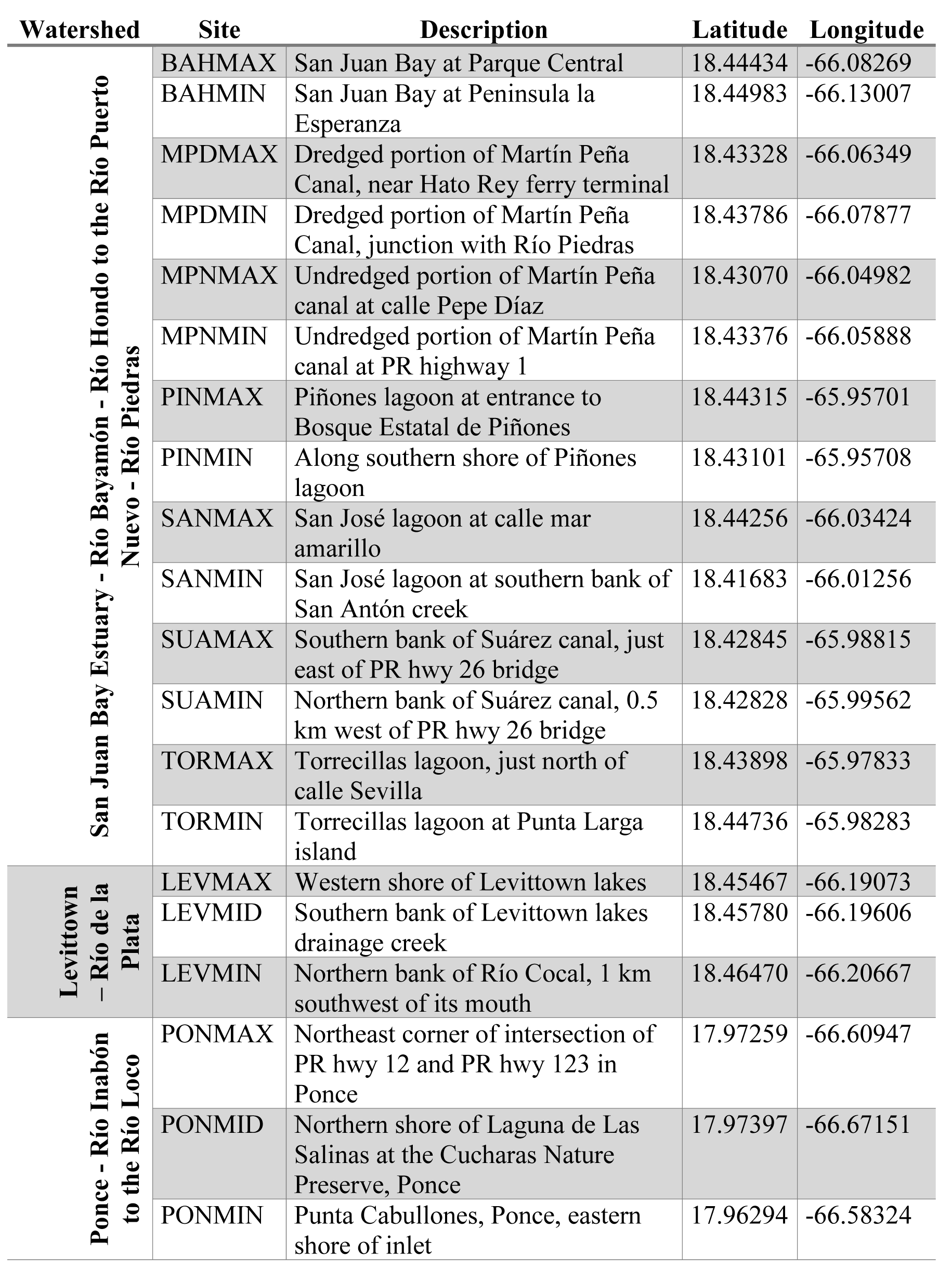
Site abbreviations used throughout the study and their corresponding locations

Roots were collected using a 3.15 cm (2.25 inch) diameter hand auger to extract soil and roots from depths of 0-10 and 10-20 cm at the same plots selected for leaf sampling. Roots were washed repeatedly with de-ionized water until no sand/dirt/silt could be seen in the rinse water, and then dried in a drying oven. Roots were then weighed for dry weight and fine roots were separated as those with a diameter less than 1mm, with coarse roots being everything else. Both leaves and roots were ground and pulverized to a powder and samples were weighed out and sent for trace metals and stable isotope analysis as described below.

Additional leaves were sampled from specimens at two herbaria in San Juan, Puerto Rico: the University of Puerto Rico’s herbarium at the Jardín Botánico Norte, and the University of Puerto Rico’s-Río Piedras herbarium at the University of Puerto Rico’s Río Piedras campus (Table 2). Specimens range in collection years from 1914 to 2011, include all mangrove species of Puerto Rico, and are from locations in San Juan and Ponce, but not Levittown. One to two leaves were taken from each specimen, also from the 2^nd^ or 3^rd^ youngest cohort, making sure to avoid leaves with glue or other apparent manipulations. No morphology measurements were taken from herbarium leaves, as they were already desiccated, and leaves were immediately pulverized in the same manner as field collected samples and sent for chemical analyses.

Leaf and root carbon and nitrogen content and stable isotope analysis was performed at the United States Environmental Protection Agency’s (EPA) Atlantic Ecology Division. Three to four milligrams of powdered tissue samples were analyzed via an Elementar Vario Micro elemental analyzer and a continuous flow Isoprime 100 isotope ratio mass spectrometer (Elementar America, Mt. Laurel, NJ). Standard reference materials USGS 40 (δ^13^C = −26.39‰; δ^15^N = −4.52‰) and USGS 41 (δ^13^C = 37.63‰; δ^15^N = 47.57‰) were also analyzed and used to normalize the isotope values of the tissue samples. Isotope values are thus expressed in reference to these standards as δ notation where δX (‰) = [(R_sample_ / R_standard_) − 1] × 103, and X is 13C or ^15^N and R is ^13^C/^12^C or ^15^N/^14^N isotopic ratio, respectively. Carbon isotope ratios from herbaria were further corrected for global atmospheric changes in ^13^C/^12^C with time as a result of anthropogenic combustion of fossil fuels (Suess 1955; Feng 1998). This change in δ^13^C of the atmosphere has been approximated as,

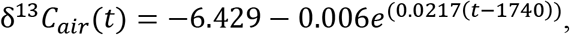

where t is the calendar year. Thus, the depletion of δ^13^𝐶𝑎𝑖𝑟 with time from the beginning of herbarium samples in 1914 is described as,

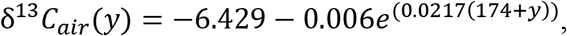

where y is the number of years since 1914. Therefore, to account for the global depletion in δ^13^C with time, and thus isolate only local changes in δ^13^C, all herbarium samples were corrected for by adding back the depleted δ^13^*C*_air_(*y*) for their collection year.

**Table 2.** Herbarium specimens from which small leaf samples were taken for chemical analyses. JBN is the herbarium at the Jardín Botanico Norte, and UPRRP is the herbarium at the University of Puerto Rico-Río Piedras. Ag is *Avicennia germinans*, Lr is *Laguncularia racemosa*, and Rm is *Rhizophora mangle*. Measurements on leaves were trace metals content (Met.) and carbon and nitrogen content and stable isotopes (C & N).

| Herbarium | ID | Sp. | Sampling Location | Collection Year | Collector | Measure |
| --- | --- | --- | --- | --- | --- | --- |
| JBN | 1875* | Lr | Martín Peña | 1914 | J.A. Stevenson | Met., C & N |
| JBN | 020912 | Rm | Martín Peña | 1914 | J.A. Stevenson | Met., C & N |
| JBN | 2189* | Lr | Martín Peña | 1914 | J.A. Stevenson | Met., C & N |
| JBN | 020913 | Rm | Martín Peña | 1914 | J.A. Stevenson | C & N |
| JBN | 032105 | Lr | Martín Peña | 1935 | Holdridge & Gerhart | Met., C & N |
| JBN | 029758 | Rm | Piñones | 1938 | L. R. Holdridge | Met., C & N |
| JBN | 032103 | Lr | Piñones | 1938 | L.R. Holdridge | Met., C & N |
| JBN | 033045 | Ag | Piñones | 1938 | L.R. Holdridge | Met., C & N |
| JBN | 032098 | Lr | Piñones | 1952 | Elbert L. Little, Jr. | C & N |
| JBN | 029763 | Rm | Piñones | 1952 | Elbert L. Little, Jr. | C & N |
| JBN | 231* | Lr | Piñones | 1968 | EGO | Met., C & N |
| JBN | 230* | Lr | Piñones | 1968 | EGO | C & N |
| UPRRP | 003335 | Rm | Piñones | 1974 | M.D. Byer | C & N |
| UPRRP | 011335 | Ag | Piñones | 1980 | M. Del Llano | C & N |
| UPRRP | 003387 | Ag | Piñones | 1984 | J. Ackerman, A. Cuevas, E. Hernández, F. Axelrod | Met., C & N |
| UPRRP | 004389 | Lr | Piñones | 1985 | F. Axelrod, J.D. & C. Turnquist, Y.A. & B Axelrod | Met., C & N |
| UPRRP | 010851 | Lr | Piñones | 1988 | S. Guzman & F. Rodriguez | C & N |
| UPRRP | 010926 | Lr | Piñones | 1988 | Brenda Molano & M. Burgos | C & N |
| UPRRP | 011698 | Lr | Piñones | 1988 | Elvia J. Melendez & Jose M. Melendez | C & N |
| UPRRP | 012094 | Rm | Piñones | 1988 | D.D. Fernández & D. García | C & N |
| UPRRP | 041374 | Lr | Piñones | 2005 | -- | C & N |
| UPRRP | 046386 | Lr | Piñones | 2005 | N.M. Martínez & M.A. Caraballo | C & N |
| UPRRP | 056582 | Rm | Piñones | 2006 | Andrew Townesmith & Greg Gust | C & N |
| UPRRP | 053344 | Ag | Martín Peña | 2009 | S. Vazquez, D. Lara & D. Ball | C & N |
| UPRRP | 053349 | Rm | Martín Peña | 2011 | W. Recart & E. Otero | Met., C & N |
| UPRRP | 053345 | Rm | Martín Peña | 2011 | W. Recart & E. Otero | C & N |
| UPRRP | 055546 | Lr | Martín Peña | 2011 | W. Recart & E. Otero | C & N |

\* Herbarium specimen IDS were illegible and an ID from the label was used instead.

Trace metals were analyzed at Florida International University’s Southeastern Environmental Research Center Environmental Analysis Research Lab. Powdered root and leaf samples were measured to roughly 0.5 g and acid digested to form a liquid matrix following EPA Method 3050B (United States Environmental Protection Agency 1996). Sample matrices were then fed through an inductively coupled plasma mass spectrophotometer to determine ppb content of silver (Ag), arsenic (As), cadmium (Cd), cobalt (Co), chromium (Cr), copper (Cu), mercury (Hg), nickel (Ni), lead (Pb), and zinc (Zn). These were converted to units of µg/g using the final volume of the matrix and the original weight of the digested tissue sample.

Surrounding land cover metrics at each site were extracted from various spatial datasets and used to characterize the urbanization of each site within a 0.5 km radius as described in more detail in (Branoff 2018b). Land cover classes of urban, vegetated, and open water were taken from the 2010 National Oceanic and Atmospheric Administration’s Coastal Change Analysis Program (C-CAP) from Puerto Rico (Office for Coastal Management 2017). Mangrove coverage was taken from a global dataset from 2012 (Hamilton and Casey 2016). Road density was taken from the 2015 United States Census Program’s TIGER line network of roads (U.S. Census Bureau 2015). Population density was taken from population totals at the block level from the 2010 United State’s Census, and corrected to assume all people lived in non-road impervious surfaces within the sampling area. These variables were used individually as predictors for measured morphology and chemistry traits, as was an urban index that includes them all and is taken as a relative level of urbanization between the sites.

Surface water chemistry measurements were taken from the San Juan Bay Estuary Program, as described in (Branoff 2018b), and are only available for San Juan sites (Figure 1). Measurements were averaged from 2012 to 2017 for each site. Monthly metrics were pH, temperature (°C), dissolved oxygen (DO) (mg/L), and salinity (PSS). Biannual metrics were total Kjeldahl nitrogen (mg/L), nitrate and nitrite total (mg/L), ammonium (mg/L), total phosphorous (mg/L), and fecal and enterococcus coliform bacterial counts (CFU/100mL). Flooding metrics were aggregated at the site level from water level models also representing the period of 2012 to 2017, and which were constructed from water-level observations at piezometers throughout the mangroves as described in (Branoff 2018b) (Figure 1). Flooding metrics include the proportion of time flooded (%), mean flood length (h), mean flood frequency(day^−1^), and mean flood depth (m).

All statistical analyses were performed in the R programming language (Yan et al. 2011).

Linear models were constructed in raw and log-transformed data of the form y~x and y~ln(x) using the *lm* function to gauge the significance of predictor variables on leaf and root morphology and chemistry. Predictor variables were those of land cover, surface water chemistry, and flooding. Additionally, leaf and root chemical measurements were used as predictor variables for morphology measurements, and trace metal content was used as a predictor of carbon and nitrogen chemistry. Analysis of variance (ANOVA) was also performed to detect differences in measured traits between species and watersheds, as done through the *aov* function (Yan et al. 2011). Scatter plots of these relationships are graphed using the ggplot function from the ggplot2 package (Wickham 2009), and linear models are shown as lines when statistically significant (p < 0.05).

## Results

### Leaf morphology and root biomass

Leaf and root morphology summaries are presented in Table 3. Leaf area was species-specific, with *Rhizophora mangle* leaves being larger, and heavier than both *Avicennia germinans* and *Laguncularia racemosa*, respectively (ANOVA; p <0.001). These differences translated to more dense leaves and a greater LMA for *Rhizophora mangle* in comparison with *Laguncularia racemosa* (ANOVA; p <0.05). All species differed in their leaf length to width ratios, with *Laguncularia racemosa* having the most rounded leaves (ratio closest to 1) and *Avicennia germinans* having the most elliptical (greatest ratio) (ANOVA; p < 0.01). In stomatal density, *Avicennia germinans* possessed 81.9 and 88.0 more stoma per mm^2^ than *Rhizophora mangle* and *Laguncularia racemosa*, respectively (ANOVA; p < 0.001). There were no differences in leaf traits between watersheds, but Levittown forests possessed 20.7 and 20.3 kg more roots per m^2^ than Ponce and San Juan, respectively (ANOVA; p < 0.05). It was not possible to assess differences in root biomass by species due to an inability to differentiate root origins and to insufficient monospecific stands.

**Table 3.**
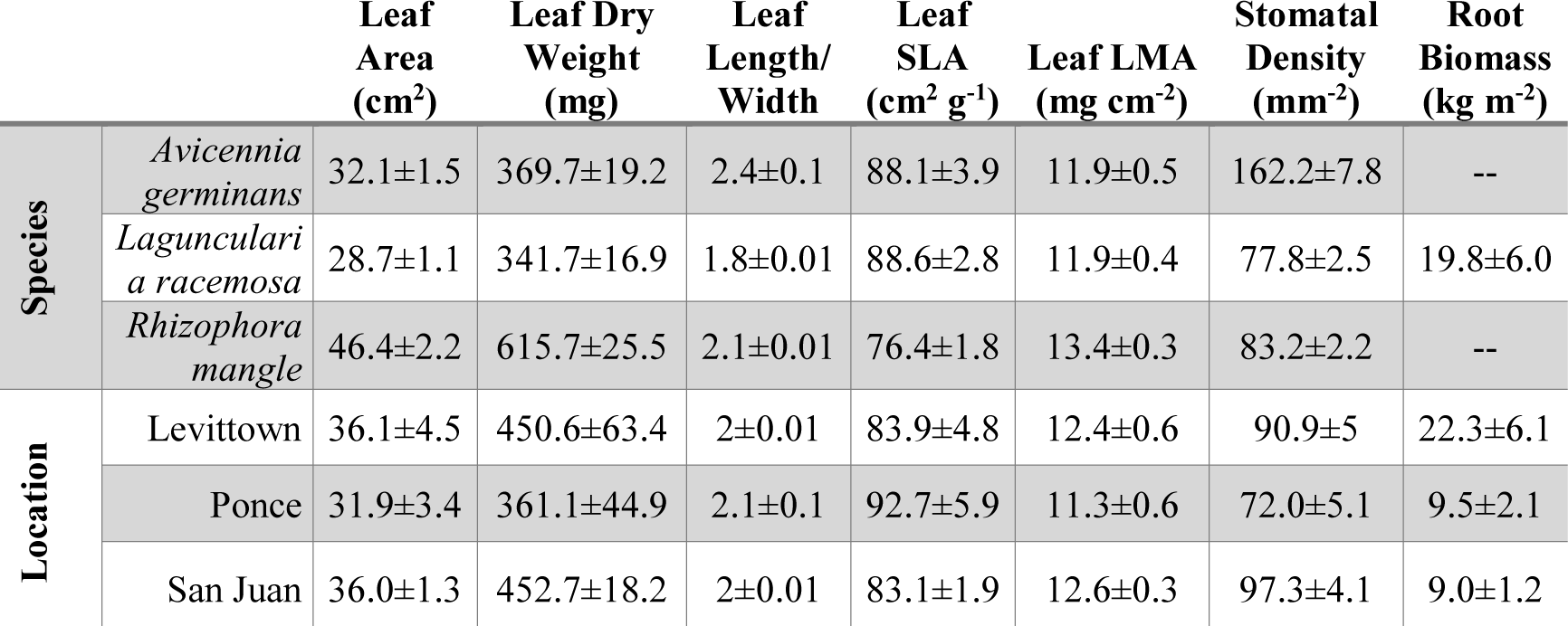
Leaf morphology and root biomass means and standard errors by species and location. Species for root measurements refer to sites in which each species represented at least 50% of the aboveground biomass.

Although some of the leaf morphological traits were correlated with urbanization, flood length, and surface water chemistry, there were no consistent patterns between species (Figure 2), while the most consistent patterns resulted from leaf chemistry, especially that of carbon and nitrogen (Figure 3). Leaf dry weight in *Avicennia germinans* was best predicted by surface water phosphorus (R^2^ = 0.35, p < 0.05). The leaf width of *Laguncularia racemosa* was best predicted by surface water salinity (R^2^ = 0.6, p < 0.01), and the SLA of *Avicennia germinans* (R^2^ = 0.61, p < 0.001) and the stomatal density of *Laguncularia racemosa* (R^2^ = 0.3, p < 0.01) were best predicted by the mean flood depth. All other morphological metrics in all species shared their strongest significant relationships with leaf chemical traits (Figure 3). Leaf area in *Laguncularia racemosa* (R^2^ = 0.43, p < 0.01) and *Rhizophora mangle* (R^2^ = 0.46, p < 0.01) was best predicted by leaf δ^13^C. Leaf dry weight in *Laguncularia racemosa* was best predicted by the leaf Ni content (R^2^ = 0.43, p < 0.01), while that in *Rhizophora mangle* was best predicted by Cd (R^2^ = 0.53, p < 0.01). For the leaf length to width ratio, the best predictor for *Laguncularia racemosa* was leaf δ^13^C (R^2^ = 0.47, p < 0.001), and that for *Avicennia germinans* was the carbon to nitrogen ratio (R^2^ = 0.40, p < 0.05). The SLA and LMA in all three species were significantly predicted by the leaf carbon to nitrogen ratio (*Avicennia germinans*; R^2^ = 0.41, p < 0.05; *Laguncularia racemosa*, R^2^ = 0.62, p < 0.001; *Rhizophora mangle*, R^2^ = 0.36, p < 0.01). Finally, the stomatal density of *Avicennia germinans* was best predicted by leaf Ni content (R^2^ = 0.89, p < 0.01), and that for *Rhizophora mangle* was best predicted by leaf δ^13^C (R^2^ = 0.3, p < 0.05).

**Figure 3.**
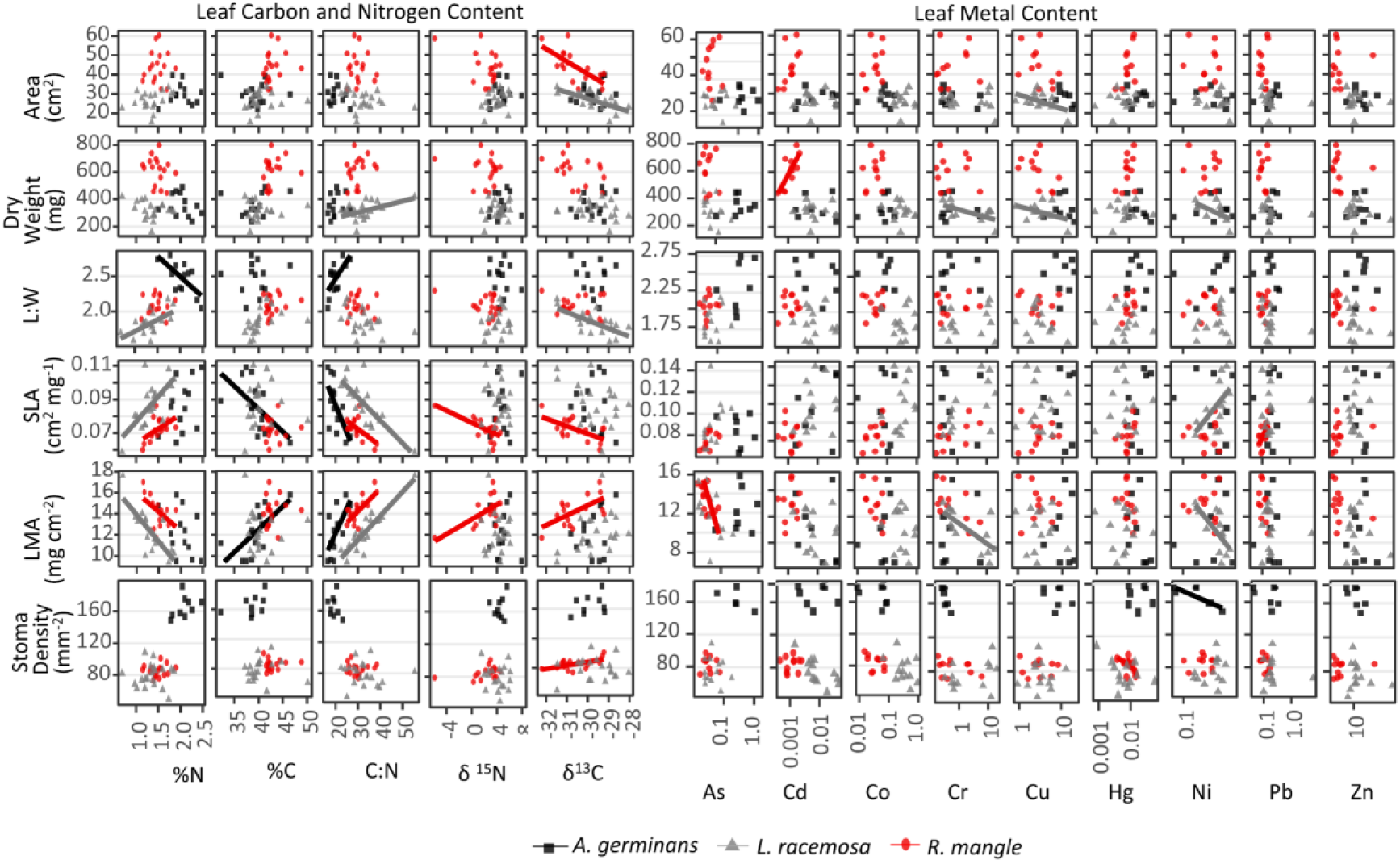
Leaf morphology traits shared their strongest and most consistent trends with leaf chemical traits, especially of carbon and nitrogen. The SLA and LMA was the only trait to be consistently tied the same predictor in all three species, in this case the carbon to nitrogen ratio.

Root biomass was not significantly correlated with urbanization or flooding metrics but did share its strongest significant relationships with root metal content and surface water chemistry. Fine root arsenic content was the strongest predictor of total root biomass (linear model; R^2^ = 0.68, p < 0.05), followed by surface water phosphorus for 0-10 cm mass (R^2^ = 0.34, p < 0.01) and root Zn content for 10-20 cm (R^2^ = 0.74, p < 0.05).

### Leaf and root chemistry

Leaf and root chemistry by species and watershed are presented in Table 4. As with morphology, leaf stable isotope and carbon and nitrogen chemistry was species, but not watershed specific. Leaf percent nitrogen was highest in *Avicennia germinans* (ANOVA; mean difference = 3.2%, p < 0.05), and the carbon to nitrogen ratio was the lowest (mean difference = 11.3, p < 0.05), but neither was different among the other species. Percent carbon was highest only in *Rhizophora mangle* (mean difference = 0.6%, p < 0.05), and both leaf δ^15^N (mean difference = 2.2‰, p < 0.01) and δ^13^C (mean difference = 0.8‰, p < 0.05) was lowest in the same species only. Trace metal contents varied across both species and watersheds. In leaves, Ag was highest at MPDMAX (0.28±0.24 µg/g), As at BAHMIN (0.44±0.22), Cd at PINMAX (0.02±0.008), Co at PONMAX (0.54±0.04), Cr at PONMAX (12.5±3.74), Cu at PINMAX (11.52±3.08), Hg at MPDMAX (0.03±0.02), Ni at PONMID (0.48±0.09), Pb at MPDMAX (2.80±2.60), and Zn at PONMID (50.76±23.30). Across all sites, leaf content of all metals in *Rhizophora mangle* were lower or similar to those of the other two species. In contrast, many of the metal levels in *Avicennia germinans* were higher than those of the other species. At all sites where species were co-inhabitants, leaves of *Avicennia germinans* held on average 321% higher content of Cu than those of *Laguncularia racemosa* and *Rhizophora mangle*. For As, at 88% of the co-inhabited sites, *Avicennia germinans* held on average 873% higher content than the other species. For Hg, Pb, and Zn, this percentage of sites with higher content in *Avicennia germinans* was 63%. For *Laguncularia racemosa*, content of Co and Cr was on average 590% higher than those of other co-inhabitants at 55% of the sites, followed by Ni and Cd, which were higher at 45% of the co-inhabited sites. For *Rhizophora mangle*, only mercury was higher than the other species, and only at 10% of the con-inhabited sites. Among watersheds, Ponce held significantly higher content of Cr, Ni and Zn in either leaves (ANOVA; p<0.05) or coarse roots (ANOVA; p<0.05) in comparison with one or both of the other watersheds.

**Table 4.** Leaf chemistry means and standard errors by species and location. Stable isotope content is in parts per thousand (‰). Metal content is in μg g^−1^ dry weight. Identification of root species’ origins was not possible, so species-specific root chemistry measurements were not obtained.

|  |  | *Avicennia*<br>*germinans* | *Laguncularia*<br>*racemosa* | *Rhizophora*<br>*mangle* | Levittown | Ponce | San Juan |
| --- | --- | --- | --- | --- | --- | --- | --- |
| $\delta^{15}\text{N}$ | Leaves | 4.7±0.3 | 4.1±0.2 | 1.8±0.4 | 2.1±1.1 | 2.4±0.5 | 3.7±0.2 |
|  | Coarse Roots | -- | -- | -- | 2.6±1.1 | 0.1±0.8 | 3.4±0.6 |
|  | Fine Roots | -- | -- | -- | 2.9±1.0 | 1.35±0.6 | 3.6±0.3 |
| $\delta^{13}\text{C}$ | Leaves | -30±0.2 | -30.1±0.1 | -30.9±0.2 | -30.9±0.3 | -30.4±0.3 | -30.3±0.1 |
|  | Coarse Roots | -- | -- | -- | -28.2±0.2 | -28.0±0.6 | -27.7±0.3 |
|  | Fine Roots | -- | -- | -- | -28.0±0.1 | -27.4±0.3 | -28.3±0.2 |
| %N | Leaves | 1.8±0.0 | 1.3±0.0 | 1.4±0.0 | 1.4±0.1 | 1.5±0.1 | 1.4±0.0 |
|  | Coarse Roots | -- | -- | -- | 0.6±0.05 | 0.7±0.18 | 0.6±0.07 |
|  | Fine Roots | -- | -- | -- | 1.1±0.08 | 1.0±0.18 | 1.3±0.05 |
| %C | Leaves | 40.4±0.6 | 40.3±0.4 | 43.6±0.3 | 43.9±0.9 | 40.6±0.5 | 41.4±0.3 |
|  | Coarse Roots | -- | -- | -- | 41.1±0.6 | 38.3±2.0 | 41.3±0.9 |
|  | Fine Roots | -- | -- | -- | 40.7±2.6 | 44.1±0.6 | 39.5±1.7 |
| C:N | Leaves | 22.4±0.6 | 34.7±1.2 | 31.5±1.7 | 31.5±1.7 | 28.6±1.2 | 31.4±0.9 |
|  | Coarse Roots | -- | -- | -- | 68.9±2.8 | 71.5±22.3 | 67.6±5.6 |
|  | Fine Roots | -- | -- | -- | 37.8±0.2 | 46.9±9.0 | 31.9±2.3 |
| Ag | Leaves | 0.034±0.016 | 0.039±0.032 | 0±0 | 0.001±0.001 | 0.012±0.012 | 0.037±0.024 |
|  | Coarse Roots | -- | -- | -- | --±-- | 0±0 | 0.2±0.1 |
|  | Fine Roots | -- | -- | -- | 0±0 | 0±-- | 0.1±0 |
| As | Leaves | 0.497±0.108 | 0.079±0.025 | 0.037±0.004 | 0.056±0.015 | 0.168±0.054 | 0.167±0.045 |
|  | Coarse Roots | -- | -- | -- | --±-- | 1.2±0.7 | 0.7±0.1 |
|  | Fine Roots | -- | -- | -- | 0.3±0.1 | 0.2±-- | 0.7±0.2 |
| Cd | Leaves | 0.017±0.004 | 0.015±0.005 | 0.001±0 | 0.003±0.001 | 0.007±0.002 | 0.013±0.004 |
|  | Coarse Roots | -- | -- | -- | --±-- | 0±0 | 0±0 |
|  | Fine Roots | -- | -- | -- | 0±0 | 0±-- | 0.1±0 |
| Co | Leaves | 0.092±0.016 | 0.403±0.06 | 0.041±0.005 | 0.339±0.097 | 0.215±0.046 | 0.177±0.041 |
|  | Coarse Roots | -- | -- | -- | --±-- | 1.8±0.7 | 2±0.8 |
|  | Fine Roots | -- | -- | -- | 2.6±0.4 | 0.7±-- | 1.3±0.3 |
| Cr | Leaves | 1.234±0.442 | 5.616±1.207 | 1.481±0.381 | 8.27±1.765 | 8.255±1.762 | 0.47±0.048 |
|  | Coarse Roots | -- | -- | -- | --N±-- | 131±117.3 | 6.7±1.6 |
|  | Fine Roots | -- | -- | -- | 8.7±7.8 | 2.1±-- | 4.4±1.2 |
| Cu | Leaves | 11.861±1.811 | 3.892±0.662 | 3.713±0.489 | 2.541±0.419 | 7.529±1.735 | 5.575±0.753 |
|  | Coarse Roots | -- | -- | -- | --±-- | 53.8±9.6 | 47.7±9.9 |
|  | Fine Roots | -- | -- | -- | 42±27.5 | 16.9±-- | 29.2±5 |
| Hg | Leaves | 0.024±0.005 | 0.013±0.006 | 0.01±0.001 | 0.009±0.002 | 0.01±0.002 | 0.017±0.004 |
|  | Coarse Roots | -- | -- | -- | --±-- | 0±0 | 0.1±0 |
|  | Fine Roots | -- | -- | -- | 0.1±0 | 0.2±-- | 0.2±0 |
| Ni | Leaves | 0.357±0.054 | 0.352±0.049 | 0.239±0.021 | 0.37±0.043 | 0.444±0.046 | 0.259±0.034 |
|  | Coarse Roots | -- | -- | -- | --N±-- | 4.8±2.1 | 2.1±0.3 |
|  | Fine Roots | -- | -- | -- | 4.7±2.9 | 1.1±-- | 2.2±0.7 |
| Pb | Leaves | 0.236±0.035 | 0.801±0.632 | 0.114±0.009 | 0.167±0.028 | 0.168±0.032 | 0.593±0.434 |
|  | Coarse Roots | -- | -- | -- | --N±-- | 25.7±3.5 | 49.6±25.7 |
|  | Fine Roots | -- | -- | -- | 15.6±3.9 | 20.1±-- | 33±2.7 |
| Zn | Leaves | 18.292±3.252 | 15.962±4.32 | 6.061±1.665 | 8.569±1.007 | 24.664±9.882 | 10.533±1.443 |
|  | Coarse Roots | -- | -- | -- | --±-- | 21.5±5.7 | 20.8±4.5 |
|  | Fine Roots | -- | -- | -- | 12.9±0.2 | 9.9±-- | 21.5±6.6 |

As with leaf morphology, relationships in leaf chemistry with urbanization were inconsistent between species and shared with surface water chemistry and leaf metal content (Figure 4). Percent nitrogen increased in *Rhizophora mangle* (R^2^ = 0.3, p < 0.05) with surface water ammonium, and in *Laguncularia racemosa* with population density (R^2^ = 0.2, p < 0.05). The carbon to nitrogen ratio decreased with every metric of urbanization in *Laguncularia racemosa* (urban index, R^2^ = 0.2, p < 0.05), and with surface water ammonium in both *Laguncularia racemosa* (R^2^ = 0.3, p < 0.05) and *Rhizophora mangle* (R^2^ = 0.3, p < 0.05). Leaf δ^15^N increased with leaf Zn content (R^2^ = 0.6, p < 0.05) in *Avicennia germinans* and decreased with surface water phosphorus in *Rhizophora mangle* (R^2^ = 0.3, p < 0.05). There were no significant relationships with leaf δ^13^C, and no relationships relating leaf carbon and nitrogen to flooding dynamics.

**Figure 4.**
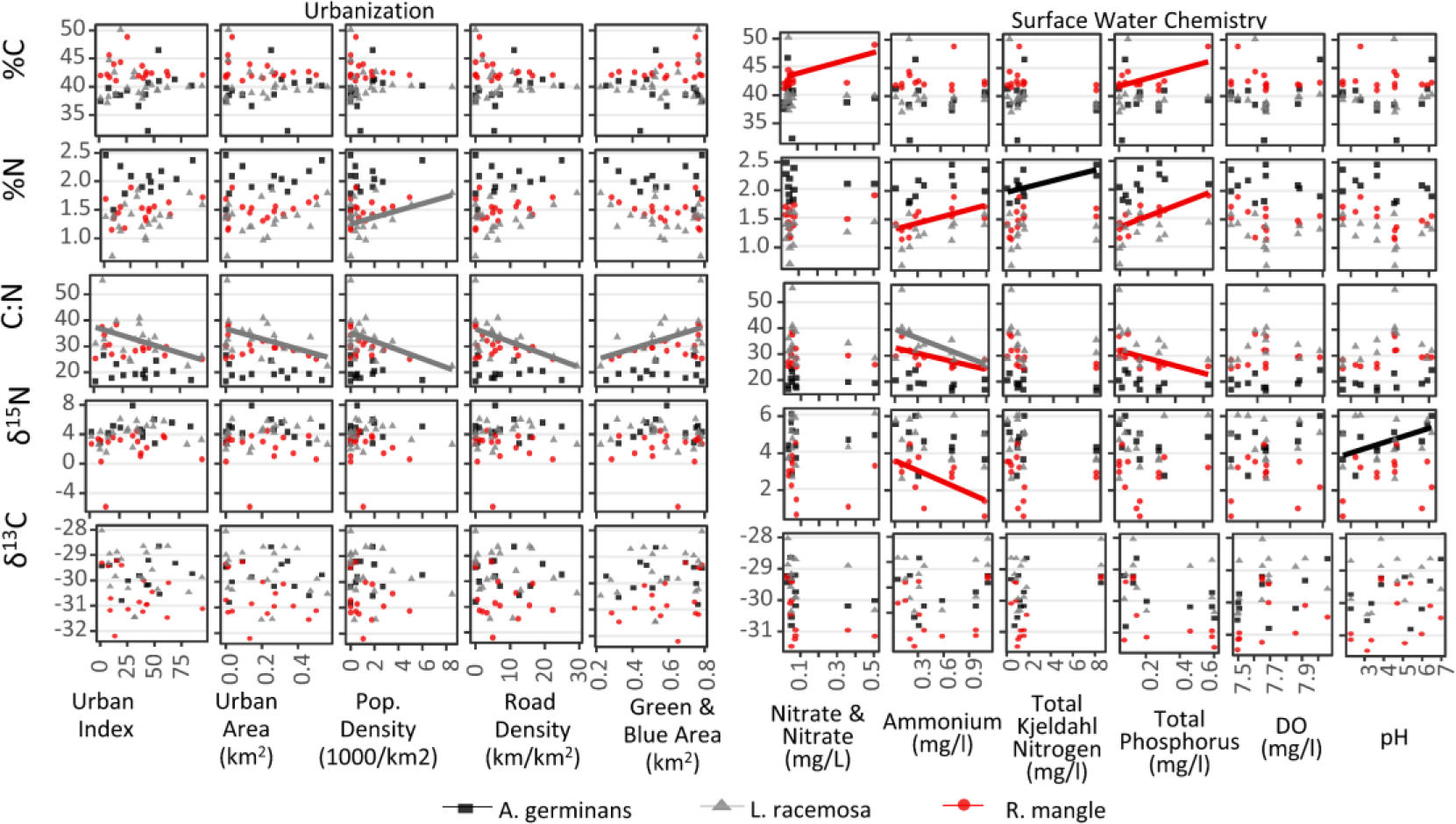
Only in *Laguncularia racemosa* was leaf carbon and nitrogen content linked to urbanization, and only for percent nitrogen and the carbon to nitrogen ratio. Otherwise, leaf carbon and nitrogen content in the other two species was best explained by surface water chemistry.

In leaf metal content, the strongest predictors were surface water chemical metrics, although some were also urbanization and flooding metrics. The number of fecal and enterococcus coliforms was the most frequent predictor, but only in *Laguncularia racemosa*, in which it was the strongest predictor of Ag (linear model; R^2^ = 0.98 p < 0.001), As (R^2^ = 0.96 p < 0.001), Cr (R^2^ = 0.51 p < 0.05), Hg (R^2^ = 0.95 p < 0.001), Pb (R^2^ = 0.97 p < 0.001), and Zn (R^2^ = 0.82 p < 0.01). Additionally, total Kjeldahl nitrogen was significantly and positively related to Cu in *Rhizophora mangle* (R^2^ = 0.88 p < 0.001) and *Laguncularia racemosa* (R^2^ = 0.60 p < 0.05), Hg in *Avicennia germinans* (R^2^ = 0.73, p < 0.05), and Zn in *Rhizophora mangle* (R^2^ = 0.88 p < 0.001). Also, content of Pb (R^2^ = 0.32, p < 0.05), Hg (R^2^ = 0.33, p < 0.05), and As (R^2^ = 0.32 p < 0.05) were all positively related to surrounding road density in *Laguncularia racemosa*.

Root carbon and nitrogen chemistry shared its strongest relationships with trace metal content, and fine and coarse roots responded differently (Figure 5). Percent nitrogen in fine roots increased with surface water Kjeldahl nitrogen (R^2^ = 0.55, p < 0.001) and to a lesser extent with surface water phosphorus (R^2^ = 0.26, p < 0.05), but all other traits responded strongest to root metal content. Percent carbon of fine roots decreased significantly with content of Co (linear model; R^2^ = 0.7, p < 0.01), Cr (R^2^ = 0.7, p < 0.001), Cu (R^2^ = 0.7, p < 0.01), and Nickel (R^2^ = 0.7, p < 0.01), while coarse root percent carbon responded positively to Cd (R^2^ = 0.4, p < 0.05) and Zn (R^2^ = 0.4, p < 0.05). Stable isotopes resulted in responses only in coarse roots, whose δ^15^N decreased with Cu (R^2^ = 0.5, p < 0.05) and Zn (R^2^ = 0.4, p < 0.05), and whose δ^13^C decreased with Zn (R^2^ = 0.6, p < 0.01). Fine root δ^13^C also decreased with the urban index (linear model; Urban Index, R^2^ = 0.5, p < 0.05), and increased with the proportion of time flooded (R^2^ = 0.3, p < 0.01). Root metal content was not consistently predicted by any of the tested variables, but Ag increased along the urban index (fine roots, R^2^ = 0.9, p < 0.001; coarse roots, R^2^ = 0.3, p < 0.05), and in fine roots, Cu increased with the proportion of time flooded (R^2^ = 0.6, p < 0.05), and both Ni (R^2^ = 0.6, p < 0.05) and As (R^2^ = 0.5, p < 0.05) increased with the flood frequency.

**Figure 5.**
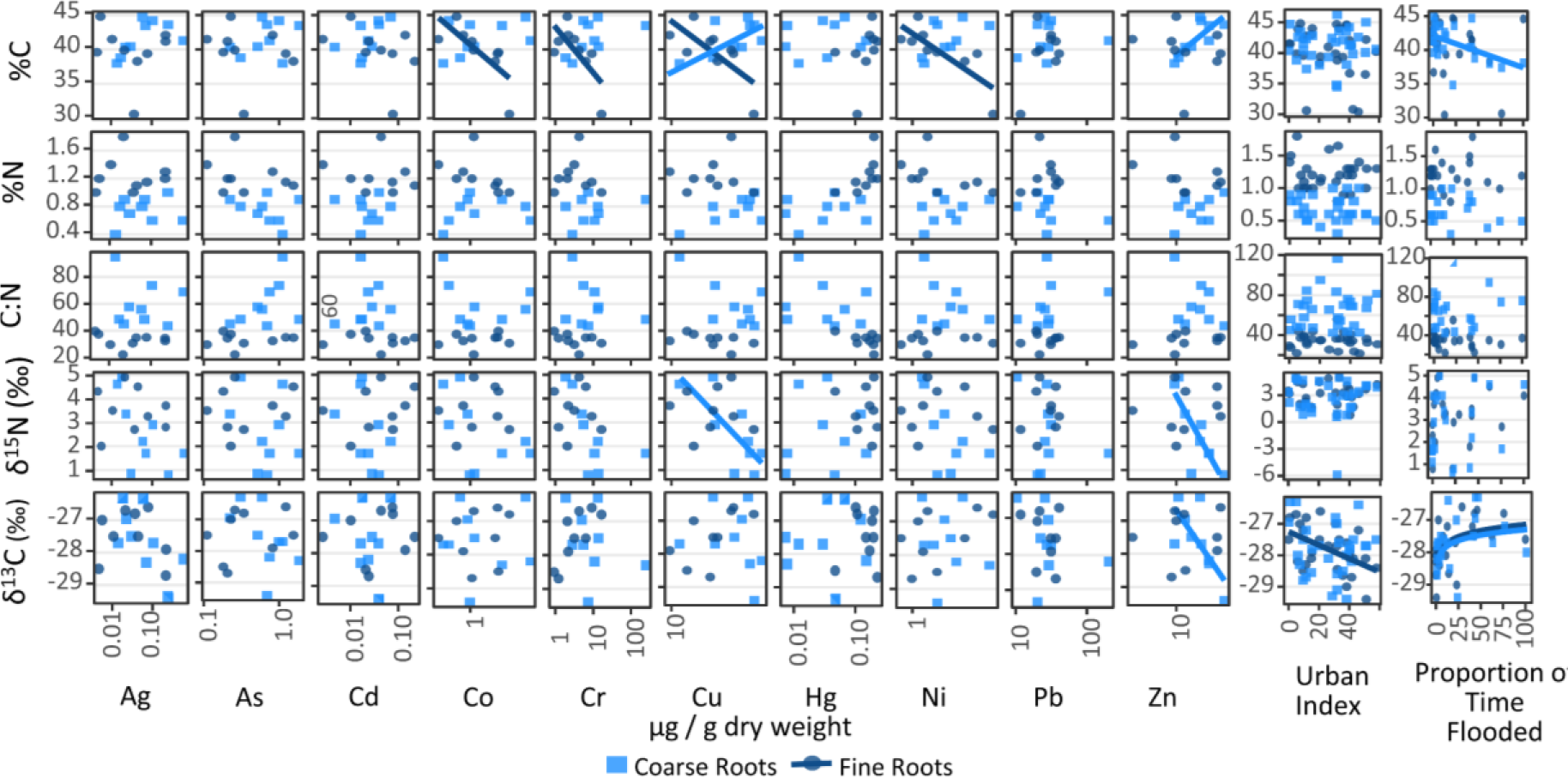
Root carbon and nitrogen chemistry was distinct between fine and coarse roots, but shared its strongest relationships with root metal content, and to a lesser extent with the urban index and the proportion of time flooded.

### Herbarium specimens

There were few consistently significant patterns in leaf chemistry with time from herbarium samples (Figure 6 & Figure 7). In Piñones, leaves of *Avicennia germinans* decreased in δ^15^N (R^2^ = 0.4, p < 0.01) and δ^13^C (R^2^ = 0.4, p < 0.01), and along with those of *Laguncularia racemosa*, increased in %N (*Avicennia germinans*, R^2^ = 0.2, p < 0.05; *Laguncularia racemosa*, R^2^ = 0.1, p < 0.05), from 1938 to 2016. In Martín Peña, leaves of *Avicennia germinans* increased in δ^15^N (R^2^ = 0.7, p < 0.001) from 2006 to 2016, and those of *Rhizophora mangle* decreased in δ^13^C (R^2^ = 0.5, p < 0.001) from 1914 to 2016. In trace metal content, there were significant patterns within species from the same water body (Figure 7). Mercury content of both *Laguncularia racemosa* and *Rhizophora mangle* from Caño Martín Peña decreased 99.99% and 99.95% from values of 88 µg/g and 102 µg/g, respectively in 1914 to 0.04 µg/g and 0.014 µg/g in 2016 (linear models; *Laguncularia racemosa*, R^2^=0.9, p < 0.05; *Rhizophora mangle*, R^2^=0.9 p < 0.01). During the same period in Caño Martín Peña, Ag content in *Laguncularia racemosa* increased 1,690% from 0.02 µg/g to 0.5 µg/g (R^2^=0.9, p < 0.05), while Zn (R^2^=0.9, p < 0.05) and Pb (R^2^=0.95, p < 0.05) from *Rhizophora Mangle* decreased 32% and 89%. In *Avicennia germinans* from Piñones, the content of Cd (R^2^=0.95, p < 0.01), Co (R^2^=0.95, p < 0.05), and Cr (R^2^=0.95, p < 0.01) decreased 53%, 83%, and 19%, respectively from 1938 to 2016. The significance of all trace metal models was maintained when non-herbarium samples from 2016 were included.

**Figure 6.**
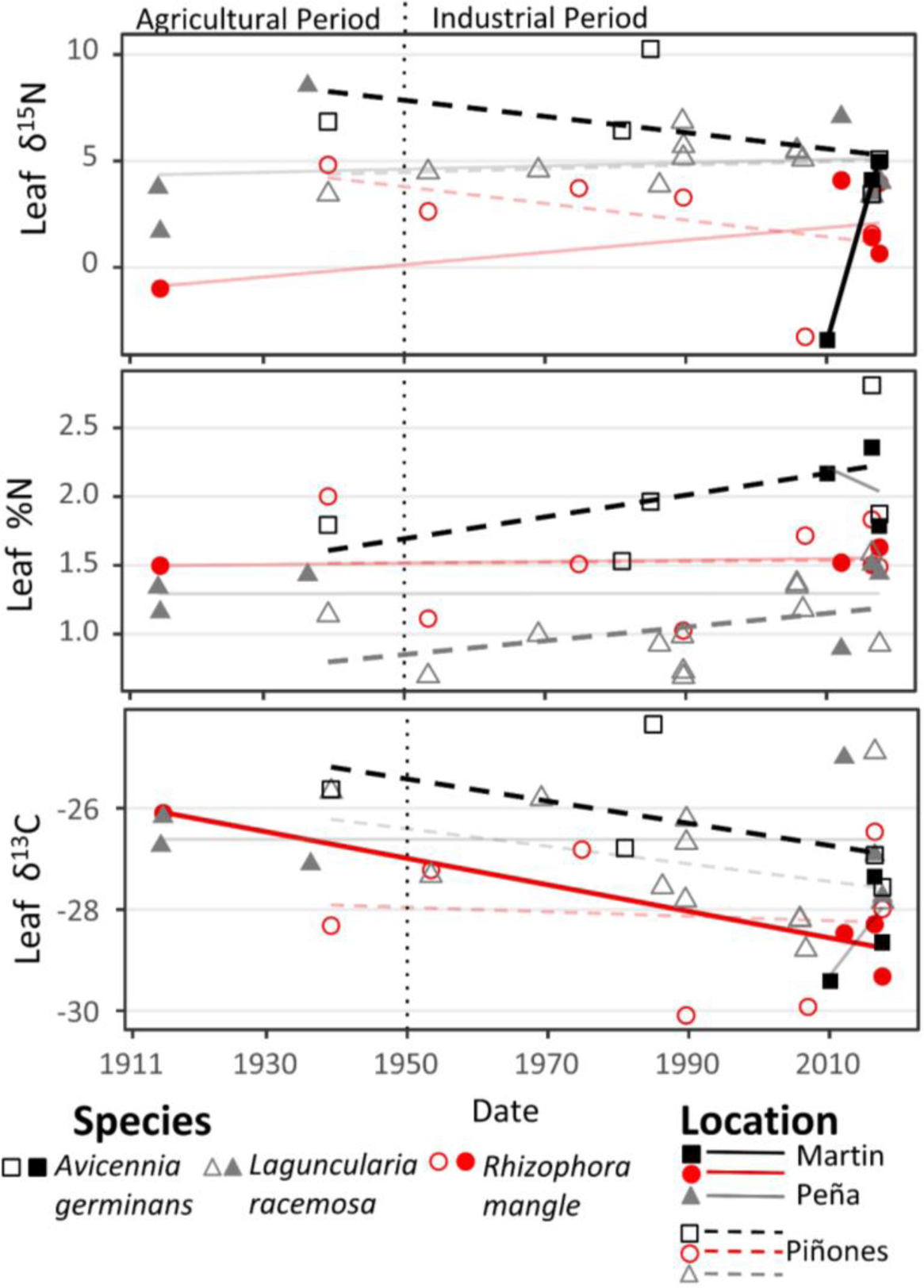
There was little evidence for any consistent change in leaf carbon and nitrogen chemistry with time from herbarium samples representing the most urban and least urban mangroves of San Juan. Bold lines are the statistically significant trends, while faded lines mark non-significant trends. Line of agricultural-industrial transition from Martinuzzi et al. (2009).

**Figure 7.**
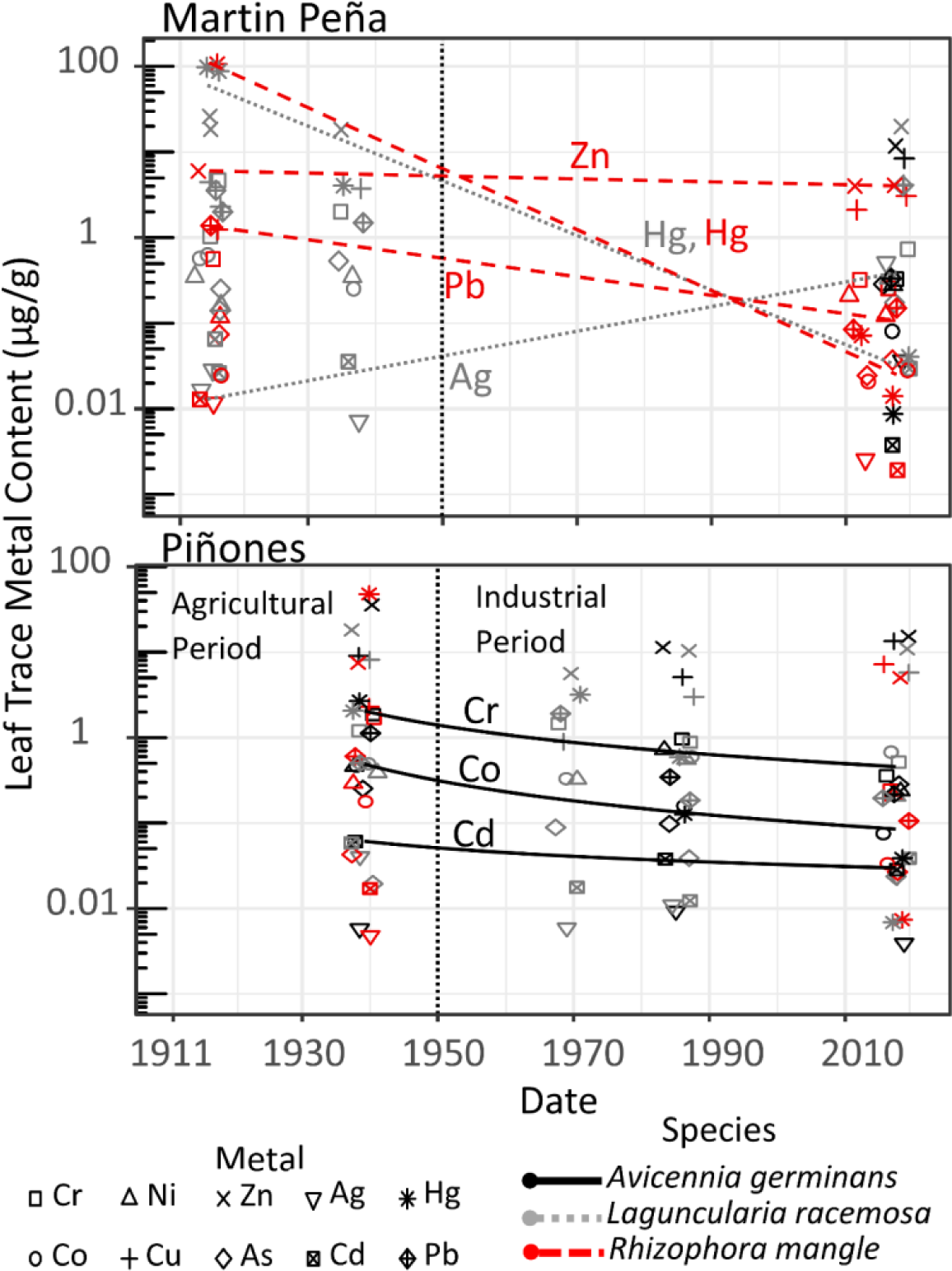
Herbarium leaf samples showed mixed changes in trace metal content with time. Mercury content decreased 99.99%, and original levels may suggest post-collection contamination of this metal. Silver content increased nearly 2000% over the last century, while Pb, Cr, Co, and Cd all decreased in one species from one location. No other metals expressed significant trends with time. Line of agricultural-industrial transition from Martinuzzi et al. (2009).

## Discussion

Plants have been shown to respond to urban environments through variation in chemical and morphological traits, and this holds true for mangroves. Yet little is known on the extent of these responses and how they compare to those of other well-known influences on mangrove traits. There was some indication from this study that urbanization is influencing mangrove leaf and root traits, but most influences were species-specific and are likely the result of secondary urban effects. These influences were only partially detected along the spatial and temporal gradients, reflecting a non-linear and non-homogenous nature of urban influences on the environment. Still, the observed patterns partially agree with previous studies on urban plant traits, as well as with studies on mangrove traits and their physiological mechanisms, as described below. For the most part, these results suggest that variation in urban mangrove habitats of Puerto Rico is manifested in both chemical and morphological traits in response to changes in water chemistry. These are likely driven by physiological processes for adjusting water-use efficiency, coping with hypoxic and toxic metal stress, and optimizing metabolism under varying nutrient availability.

Some of the strongest and most consistent links to urbanization involved nitrogen and phosphorus (Figure 4). Percent nitrogen of roots as well as leaves in all three species increased with either nitrogen, phosphorus, or population density. This agrees with other studies on both mangroves and terrestrial plants and is hypothesized to be a response to nutrient enrichment in urban environments sourced from both sewage and roads (Kendall et al. 2007; Branoff 2017b). But some of the remaining chemical traits did not support this hypothesis. Nitrogen sourced from human waste and roads should be enriched in δ^15^N (Savage 2005; Kendall et al. 2007), and mangroves will increase foliar δ^15^N when enriched with phosphorus or with nitrogen from sewage (Feller et al. 2003; Ernesto Medina et al. 2010). But field collected leaves showed no trend in δ^15^N with urbanization, and herbarium leaves showed no consistent increase in nitrogen content, despite a significant increase in San Juan’s population and an island wide transition from an agricultural to industrial economy over the range of dates covering herbarium specimen collections. Therefore, although both nitrogen and phosphorus are enriched in urban trees, or trees where water nitrogen and phosphorus is elevated, the source of the nitrogen does not seem to be sewage.

Fry et al. (2000) offer two potential models for observed variations in δ^15^N of anthropogenically influenced mangroves in south Florida. Model one suggests that baseline δ^15^N is high, and observed enrichment is due primarily to in-plant fractionation and resulting δ^15^N depletion occurs as a result of differences in available nitrogen types (ammonium, nitrate, nitrite etc.) and quantities relative to demand. Model two suggests that baseline δ^15^N is low, and that higher microbial fractionation and nitrogen losses between the nitrogen source and the plants as a result of nitrogen enrichment results in higher δ^15^N in mangroves (Reis, Nardoto, Rochelle, et al. 2017; Reis, Nardoto, and Oliveira 2017). The authors reject model two because their observed minimal δ^15^N of −5 ‰ is too low to be a baseline, and because they observed no increase in δ^15^N in more productive systems. In contrast, the present study reflects what is probably mostly representative of model two.

The minimum of site means for observed foliar δ^15^N was −1.4 ‰, sufficiently high to be a baseline nitrogen sourced from marine and fixation inputs (Carpenter et al. 1997; Fry et al. 2000). Further, foliar δ^15^N increased with mangrove stand biomass, as predicted by model two (Figure 8). Nearly all site means in foliar δ^15^N were higher than that of PONMIN (1.4‰), which is the least urban of the open marine sites and thus most reflective of the marine baseline values. This suggests that baseline nitrogen is primarily marine, with a δ^15^N value of 1.4‰ and that increases in δ^15^N result from higher nitrogen availability and the resulting increase in nitrification, denitrification, and nitrogen loss rates between the source nitrogen and the mangroves. But if the added nitrogen is sewage, δ^15^N values would be expected to be even higher (~8‰) (Savage 2005), suggesting there is an additional source of nitrogen enrichment in these systems.

**Figure 8.**
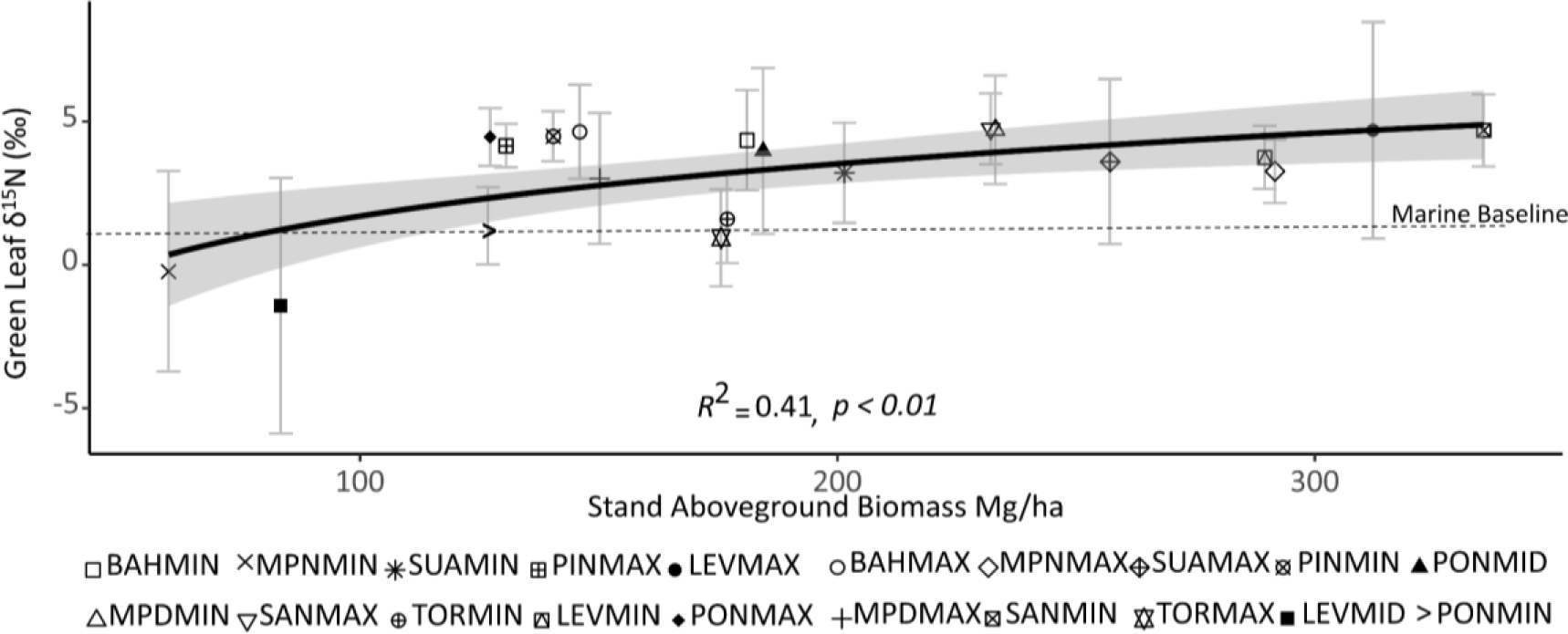
Foliar **δ^15^N** increase with stand biomass, as taken from Branoff & Martinuzzi, 2018. This suggests variations in nitrogen stable isotopes are due not to source variability, but to overall site fertility and fractionation processes between the source and the mangroves.

Based on previous studies of mangroves and on terrestrial urban flora, additional nitrogen sources may be either roads, nitrogen fixation, or both. Nitrogen fixation has been shown many times in mangroves, and has even been estimated to supply up to 60% of nitrogen in some systems (Zuberer and Silver 1978; Paerl et al. 1981; Van der Valk and Attiwill 1984; Romero et al. 2012). Further, in many coastal systems phosphorus is a co-limiting nutrient (Howarth and Marino 2006; Bracken et al. 2015), and phosphorus enrichment stimulates this fixating community (Romero et al. 2012). Thus, with sewage derived ammonium and phosphorus, urban mangroves augment their nitrogen content (Feller et al. 1999), but their δ^15^N remains low due to a pool of nitrogen provided by fixation. This is especially true for long-term phosphorus fertilization (Romero et al. 2012), such as is the case for the mangroves of San Juan, which have been subjected to raw sewage inputs for decades (Hunter and Arbona 1995). It’s likely this relationship is not directly tied to the urbanization variables because point-source sewage inputs only influence the nearest mangrove sites. This was evident in the heterogenous surface water nitrogen and phosphorus concentrations from (Branoff 2018b).

Roads are another potential explanation for the relatively depleted δ^15^N of mangrove leaves compared to those of other urban mangroves. Previous urban herbarium analyses have shown depletion of foliar δ^15^N with time due to increasing atmospheric nitrate-nitrogen deposition from roads (Huang et al. 2016), which may further counteract any δ^15^N enrichment from sewage. But the present study found no relationship between δ^15^N and surrounding road density, suggesting either minimal influence of roads, or confounding influence of roads, fixation, and the ocean.

The same models proposed by Fry et al. (2000), in combination with the known history of the San Juan Bay Estuary, could also explain the historical trends in nitrogen content from herbarium samples. The Martín Peña canal has a history of receiving un-treated sewage from unregulated residential development, as well as sediments from various dredging campaigns throughout its development history (Environmental Protection Agency 2007; United States Army Corps of Engineers 2015). This area of the city was already significantly developed at the turn of the century, with a population density of around 500 people/km^2^, and was undergoing rapid population growth at the time of the first herbarium measurements in 1914 (50% in Martín Peña bordering neighborhoods between 1910 and 1920) (United States Census Bureau 1913). This may be why leaf δ^15^N and %N remained unchanged over the next century, because they were already elevated from sewage exposure at the time of the first herbarium specimen collection. At Piñones, however, population density from the same census was 50 people/km^2^ and population growth was 2%, numbers that do not support the relatively low nitrogen and high δ^15^N content of leaves from this time. Instead, these values may be explained by a switch in the dominant nitrogen form, from nitrate to ammonium. Prior to the 1950’s the Piñones lagoon likely received water through a series of drainage and navigation canals from the Río Grande de Loíza (Webb and Gómez-Gómez 1998; Cordero 2015), which is the largest river by volume on the island and which then drained agricultural lands throughout its basin. The nitrogen of this likely nitrate rich agricultural water would have undergone elevated microbial fractionation, leaving a pool of enriched δ^15^N for the mangroves (model 1, (Fry et al. 2000)). Following the damming of the Río Grande de Lóiza in 1953, as well as the agricultural-industrial transition at the same time, the Piñones lagoon likely switched from a nitrate to an ammonium system. Ammonium does not fractionate as much as nitrate, so leaf %N increased and δ^15^N decreased as population density increased to around 2000 people/km^2^ in the year 2000 (United States Census Bureau), and presumably sewage sourced nitrogen and phosphorus inputs also increased.

Sewage inputs, along with changes in environmental regulation and water treatment infrastructure, may also explain mangrove metal content from both herbarium and modern samples. Metal content in herbarium leaves was either static or decreased with time, in agreement with sediment records from Webb and Gómez-Gómez (1998), and likely a result of improvements to the waste and storm water infrastructure in the estuary, especially after its inclusion in the national estuary program in 1993. The most dramatic change in metal content was that of mercury from Martín Peña. The Hg content of these leaves in 1914 (~100 µg/g) was roughly fifty times greater than the maximum reported from other systems (Bayen 2012), and 500 times greater than field collected leaves from the same site. This may be the true leaf content of these samples, but it may also be a result of the early use of mercury chloride as a pesticide in herbaria, which could not be confirmed from the herbaria in this study. Still, the modern mangroves in Martín Peña, which has the longest history of urbanization from this study, harbor some of the highest metal levels compared to the other forests. The MPDMAX site at Martín Peña registered as the highest for more metals in various tissues than any other site, and the only site to register the highest in at least one metal for all three tissue compartments (leaves, coarse roots, and fine roots). These metals were Ag in all tissues (leaves 0.275±0.238 µg/g, coarse roots 0.318±0.147 µg/g, and fine roots 0.217±0.001 µg/g), Cd (0.034±0.005 μg/g), Co (2.304±1.544 µg/g), and Cu (60.771±16.823 μg/g) in coarse roots, Hg in all tissues (leaves 0.032±0.023 μg/g, coarse roots 0.076±0.038 µg/g, and fine roots 0.201±0.029 µg/g), Pb in leaves (2.796±2.596 μg/g) and coarse roots (84.935±59.173 μg/g), and Zinc in coarse roots (30.527±6.616 μg/g). With the exception of Ag, all metals have either remained unchanged or decreased with time in the mangroves of Puerto Rico, and modern metal content ranges from low to moderate in comparison with those of the most commonly studied metals from other mangroves (Bayen 2012). But the content of these metals in *Laguncularia racemosa*, which is the dominant species in the most urban forests, was positively related to surface water fecal and enterococcus coliform counts, suggesting a link to untreated sewage.

Although low to moderate in comparison with other studies, the metal content in mangrove tissues may by imparting a physiological response as indicated in the other traits. Leaf stomatal densities and fine root carbon and nitrogen traits, and to a lesser extent those of coarse roots, shared their strongest relationships with the content of various trace metals. Accumulations of metals in fine roots have been noted in other studies, and are hypothesized to be a strategy for building tolerance to these toxic elements (Gussarsson 1994; Weis and Weis 2004). Also, arsenic was the strongest predictor of root biomass, which decreased as this metal content increased. The effect of this metal on mangrove roots has been noted in another study (Da Souza et al. 2014), which also notes the adaptive capacity of *Laguncularia racemosa* to mitigate the toxicity of varying anthropogenic contaminants, including metals. Indeed, this species showed the greatest number of morphological responses to metal content in the present study, and its dominance over the other species increases in urban areas, where it often forms monospecific stands (Branoff and Martinuzzi 2018). This suggests *Laguncularia racemosa* may be particularly capable of inhabiting urban landscapes, due to an ability to cope with toxic contaminants through morphological variations. But these same variations could also be predicted by leaf chemistry, which may be indirectly influenced by urbanization as well.

Leaf SLA and LMA were the most consistent traits in terms of relationships among all three species, with each species responding to at least one of the metrics of carbon and nitrogen content. Both have been tied to net photosynthetic capacity and/or to internal CO2 resistance (P B Reich et al. 1998; Hultine and Marshall 2000), which suggests a potential effect on productivity.

But these traits were also predicted by surface water chemistry and leaf metal content, inferring there may be both compounding and conflicting influences from the various environmental components. Further, mangroves have been shown to vary both morphological and chemical leaf traits in response to non-anthropogenic influences (Lin and Sternberg 1992; E Medina and Francisco 1997; Ernesto Medina 1999), and aside from stomatal density, there were no strong morphological relationships in the present study with urbanization, suggesting the observed trends in leaf morphology are due primarily to non-urban related variations in the environment. Similarly, although variations in root δ^13^C could be predicted by the urban index, they could also be predicted by the proportion of time flooded, suggesting this trait may be a response to the accumulation of various carbon compounds during hypoxic metabolism (McKee and Mendelssohn 1987) and not to the urban environment.

Roads are another potential source of toxic metals that may influence plant morphology (Lichtfouse et al. 2003; Honour et al. 2009; Bettez et al. 2013). There was some evidence for this in the observed mangrove traits of stomatal density and metal content. A number of metals increased in both root and foliar content with road density, supporting the hypothesis that roads are a major source of trace metals for surrounding ecosystems (Tomašević et al. 2005; De Nicola et al. 2008). Similarly, although only in *Rhizophora mangle*, the stomatal density decreased with every metric of urbanization, including road density. This is in agreement with one study (Verma and Singh 2006), but opposite of that observed in another species (Kardel et al. 2010), showing that species-specific responses to the urban environment may not always agree. This may be why only *Rhizophora mangle* responded in stomatal density, because this particular trait is more sensitive in this species. Also, the above mentioned influence of atmospheric nitrogen deposition sourced from roads my also explain the lower than expected foliar δ^15^N. Still, there was no observed decrease in foliar δ^13^C with any metric of urbanization, which has also been shown to occur in response to higher assimilation of fossil fuel sourced carbon near roads and highways (Lichtfouse et al. 2003).

The lack of evidence for any link between δ^13^C and urbanization as well as any reduced belowground biomass allocation from urban nitrogen enrichment (Gift et al. 2010), is probably because these traits in mangroves are also tied to plant responses to variable flooding and salinity (Lin and Sternberg 1992; E Medina and Francisco 1997; Castañeda-Moya et al. 2011). Isolating the various influences of the environment on mangrove eco-physiology is made difficult by the dynamic nature of these forests. Variations in tidal inundation, salinity and nutrient levels are well documented controls on these systems, each of which acts in both isolation and in synchrony with the others to control mangrove biology. Urban environments further complicate these dynamics by introducing novel conditions that again act in both isolation and synchrony with the other controls. This study showed that some mangrove leaf and root traits can be explained by the urban environment, especially by the presence of elevated nutrients and trace metals in urban waters. But truly isolating these from other well-known influences on mangrove eco-physiology will require well designed experiments. Such research will be under increasingly high demand as mangroves worldwide face ongoing urbanization, and as managers seek to understand how this disturbance influences mangrove function and the provisioning of ecosystem services.

## References

Balasooriya, B L W K, R Samson, F Mbikwa, U W A Vitharana, P Boeckx, and M Van Meirvenne. 2009. Biomonitoring of urban habitat quality by anatomical and chemical leaf characteristics. Environmental and Experimental Botany 65: 386–394.doi:http://dx.doi.org/10.1016/j.envexpbot.2008.11.009.

Barceló, JUAN, and Ch Poschenrieder. 1990. Plant water relations as affected by heavy metal stress: a review. Journal of plant nutrition 13. Taylor & Francis: 1–37.

Bardgett, Richard D, Liesje Mommer, and Franciska T De Vries. 2014. Going underground: root traits as drivers of ecosystem processes. Trends in Ecology & Evolution 29. Elsevier: 692–699.

Basyuni, M, D A Keliat, M U Lubis, N B Manalu, A Syuhada, and R Wati. 2018. Growth and root development of four mangrove seedlings under varying salinity. In IOP Conference Series: Earth and Environmental Science, 130:12027. IOP Publishing.

Bayen, Stéphane. 2012. Occurrence, bioavailability and toxic effects of trace metals and organic contaminants in mangrove ecosystems: a review. Environment international 48: 84–101.

Bettez, Neil D, Roxanne Marino, Robert W Howarth, and Eric A Davidson. 2013. Roads as nitrogen deposition hot spots. Biogeochemistry 114. Springer: 149–163.

Bini, Claudio, Mohammad Wahsha, Silvia Fontana, and Laura Maleci. 2012. Effects of heavy metals on morphological characteristics of Taraxacum officinale Web growing on mine soils in NE Italy. Journal of Geochemical Exploration 123. Elsevier: 101–108.

Bracken, Matthew E S, Helmut Hillebrand, Elizabeth T Borer, Eric W Seabloom, Just Cebrian, Elsa E Cleland, James J Elser, Daniel S Gruner, W Stanley Harpole, and Jacqueline T Ngai. 2015. Signatures of nutrient limitation and co‐limitation: responses of autotroph internal nutrient concentrations to nitrogen and phosphorus additions. Oikos 124. Wiley Online Library: 113–121.

Branoff, Benjamin Lee. 2017a. Quantifying the influence of urban land-use on mangrove biology and ecology: A meta-analysis. Global Ecology & Biogeography 26: 1339–1356.

Branoff, Benjamin Lee. 2017b. Quantifying the influence of urban land use on mangrove biology and ecology: A meta-analysis. Global Ecology and Biogeography 26. doi:10.1111/geb.12638.

Branoff, Benjamin Lee. 2018a. Urban Mangrove Biology and Ecology: Emergent Patterns and Management Implications. In Threats to Mangrove Forests, 521–537. Springer.

Branoff, Benjamin Lee. 2018b. Urbanization plays a minor role in the flooding and surface water chemistry of Puerto Rico’s mangroves. bioRxiv. doi:10.1101/423434.

Branoff, Benjamin Lee, and Sebastian Martinuzzi. 2018. Mangrove forest structure and composition along an urban gradient in Puerto Rico. In prep.

Carpenter, Edward J, H Rodger Harvey, Brian Fry, and Douglas G Capone. 1997. Biogeochemical tracers of the marine cyanobacterium Trichodesmium. Deep Sea Research Part I: Oceanographic Research Papers 44. Elsevier: 27–38.

Carreras, Hebe A, Martha S Cañas, and María L Pignata. 1996. Differences in responses to urban air pollutants by Ligustrum lucidum Ait. and Ligustrum lucidum Ait. f. tricolor (Rehd.) Rehd. Environmental Pollution 93. Elsevier: 211–218.

Castañeda-Moya, Edward, Robert R Twilley, Victor H Rivera-Monroy, Brian D Marx, Carlos Coronado-Molina, and Sharon M L Ewe. 2011. Patterns of root dynamics in mangrove forests along environmental gradients in the Florida Coastal Everglades, USA. Ecosystems 14. Springer: 1178–1195.

Cintrón, Gilberto, Ariel E Lugo, Douglas J Pool, and Greg Morris. 1978. Mangroves of arid environments in Puerto Rico and adjacent islands. Biotropica. JSTOR: 110–121.

Cordero, Juan Giusti. 2015. Trabajo y vida en el mangle: “Madera negra” y carbón en Piñones (Loíza), Puerto Rico (1880-1950). Caribbean Studies. JSTOR: 3–71.

Defew, Lindsey H, James M Mair, and Hector M Guzman. 2005. An assessment of metal contamination in mangrove sediments and leaves from Punta Mala Bay, Pacific Panama. Marine pollution bulletin 50: 547–552.

Diaz, S, J G Hodgson, K Thompson, M Cabido, J H C 3 Cornelissen, A Jalili, G Montserrat‐ Marti, J P Grime, F Zarrinkamar, and Y Asri. 2004. The plant traits that drive ecosystems: evidence from three continents. Journal of vegetation science 15. Wiley Online Library: 295–304.

Dolan, Rebecca W, Marcia E Moore, and Jessica D Stephens. 2011. Documenting effects of urbanization on flora using herbarium records. Journal of Ecology 99. Wiley Online Library: 1055–1062.

Domínguez, María T, Cristina Aponte, Ignacio M Pérez-Ramos, Luis V García, Rafael Villar, and Teodoro Marañón. 2012. Relationships between leaf morphological traits, nutrient concentrations and isotopic signatures for Mediterranean woody plant species and communities. Plant and Soil 357. Springer: 407–424.

Environmental Protection Agency. 2007. National estuary program coastal condition report, chap 7: Puerto Rico: San Juan Bay estuary partnership coastal condition.

Feller, Ilka C, Karen L McKee, Dennis F Whigham, and John P O’neill. 2003. Nitrogen vs. phosphorus limitation across an ecotonal gradient in a mangrove forest. Biogeochemistry 62. Springer: 145–175.

Feller, Ilka C, Dennis F Whigham, John P O’Neill, and Karen L McKee. 1999. Effects of nutrient enrichment on within‐stand cycling in a mangrove forest. Ecology 80. Wiley Online Library: 2193–2205.

Feng, Xiahong. 1998. Long-term c i/c a response of trees in western North America to atmospheric CO 2 concentration derived from carbon isotope chronologies. Oecologia 117. Springer: 19–25.

Flores-de-Santiago, Francisco, John M Kovacs, and Francisco Flores-Verdugo. 2012. Seasonal changes in leaf chlorophyll a content and morphology in a sub-tropical mangrove forest of the Mexican Pacific. Marine Ecology Progress Series 444: 57–68.

Fry, B, A L Bern, M S Ross, and J F Meeder. 2000. δ15N studies of nitrogen use by the red mangrove, Rhizophora mangle L. in south Florida. Estuarine, Coastal and Shelf Science 50. Elsevier: 291–296.

Gift, Danielle M, Peter M Groffman, Sujay S Kaushal, and Paul M Mayer. 2010. Denitrification potential, root biomass, and organic matter in degraded and restored urban riparian zones. Restoration Ecology 18. Wiley Online Library: 113–120.

Gussarsson, Monika. 1994. Cadmium‐induced alterations in nutrient composition and growth of betula pendula seedlings: The significance of fine roots as a primary target for cadmium toxicity. Journal of Plant Nutrition 17. Taylor & Francis: 2151–2163.

Hamilton, Stuart, and Daniel Casey. 2016. Creation of a high spatiotemporal resolution global database of continuous mangrove forest cover for the 21st Century (CGMFC-21). Global Ecology & Biogeography 25.6: 729–738.

Honour, Sarah L, J Nigel B Bell, Trevor W Ashenden, J Neil Cape, and Sally A Power. 2009. Responses of herbaceous plants to urban air pollution: Effects on growth, phenology and leaf surface characteristics. Environmental Pollution 157: 1279–1286. doi:https://doi.org/10.1016/j.envpol.2008.11.049.

Howarth, Robert W, and Roxanne Marino. 2006. Nitrogen as the limiting nutrient for eutrophication in coastal marine ecosystems: evolving views over three decades. Limnology and Oceanography 51. Wiley Online Library: 364–376.

Huang, Zhiqun, Bao Liu, Murray Davis, Jordi Sardans, Josep Peñuelas, and Sharon Billings. 2016. Long‐term nitrogen deposition linked to reduced water use efficiency in forests with low phosphorus availability. New Phytologist 210. Wiley Online Library: 431–442.

Hultine, K R, and J D Marshall. 2000. Altitude trends in conifer leaf morphology and stable carbon isotope composition. Oecologia 123. Springer: 32–40.

Hunter, John M, and Sonia I Arbona. 1995. Paradise lost: an introduction to the geography of water pollution in Puerto Rico. Social Science & Medicine 40. Citeseer: 1331–1355.

Kardel, Fatemeh, Karen Wuyts, M Babanezhad, Tatiana Wuytack, Geert Potters, and Roeland Samson. 2010. Assessing urban habitat quality based on specific leaf area and stomatal characteristics of Plantago lanceolata L. Environmental Pollution 158. Elsevier: 788–794.

Kendall, Carol, Emily M Elliott, and Scott D Wankel. 2007. Tracing anthropogenic inputs of nitrogen to ecosystems. Stable isotopes in ecology and environmental science. Wiley Online Library: 375–449.

Lichtfouse, Éric, Michel Lichtfouse, and Anne Jaffrezic. 2003. δ 13 C Values of Grasses as a Novel Indicator of Pollution by Fossil-Fuel-Derived Greenhouse Gas CO2 in Urban Areas. Environmental Science and Technology 37: 87–89.

Lima, Josanidia Santana, E B Fernandes, and W N Fawcett. 2000. Mangifera indica and Phaseolus vulgaris in the bioindication of air pollution in Bahia, Brazil. Ecotoxicology and Environmental Safety 46. Elsevier: 275–278.

Lin, G H, and L da S L Sternberg. 1992. Effect of growth form, salinity, nutrient and sulfide on photosynthesis, carbon isotope discrimination and growth of red mangrove (Rhizophora mangle L.). Functional Plant Biology 19. CSIRO: 509–517.

Lira-Medeiros, Catarina Fonseca, Christian Parisod, Ricardo Avancini Fernandes, Camila Souza Mata, Monica Aires Cardoso, and Paulo Cavalcanti Gomes Ferreira. 2010. Epigenetic variation in mangrove plants occurring in contrasting natural environment. PLoS One 5. Public Library of Science: e10326.

MacFarlane, G R. 2003. Chlorophyll a fluorescence as a potential biomarker of zinc stress in the grey mangrove, Avicennia marina (Forsk.) Vierh. Bulletin of environmental contamination and toxicology 70: 90–96.

MacFarlane, G R, and M D Burchett. 2001. Photosynthetic pigments and peroxidase activity as indicators of heavy metal stress in the grey mangrove, Avicennia marina (Forsk.) Vierh. Marine pollution bulletin 42: 233–240.

MacFarlane, G R, and M D Burchett. 2002. Toxicity, growth and accumulation relationships of copper, lead and zinc in the grey mangrove Avicennia marina (Forsk.) Vierh. Marine Environmental Research 54. Elsevier: 65–84.

MacFarlane, Geoff R, Claudia E Koller, and Simon P Blomberg. 2007. Accumulation and partitioning of heavy metals in mangroves: A synthesis of field-based studies. Chemosphere 69: 1454–1464. doi:http://dx.doi.org/10.1016/j.chemosphere.2007.04.059.

Martín, J A Rodríguez, C De Arana, J J Ramos-Miras, C Gil, and R Boluda. 2015. Impact of 70 years urban growth associated with heavy metal pollution. Environmental Pollution 196. Elsevier: 156–163.

Martinuzzi, Sebastian, William A Gould, Ariel E Lugo, and Ernesto Medina. 2009. Conversion and recovery of Puerto Rican mangroves: 200 years of change. Forest Ecology and Management 257: 75–84.

McKee, Karen L. 1996. Growth and physiological responses of neotropical mangrove seedlings to root zone hypoxia. Tree physiology 16. Heron Publishing: 883–889.

McKee, Karen L, Ilka C Feller, Marianne Popp, and Wolfgang Wanek. 2002. Mangrove isotopic (δ15N and δ13C) fractionation across a nitrogen vs. phosphorus limitation gradient. Ecology 83. Wiley Online Library: 1065–1075.

McKee, Karen L, and Irving A Mendelssohn. 1987. Root metabolism in the black mangrove (Avicennia germinans (L.) L): response to hypoxia. Environmental and Experimental Botany 27. Elsevier: 147–156.

Medina, E, and M Francisco. 1997. Osmolality and ð” C of Leaf Tissues of Mangrove. Estuarine, Coastal and Shelf Science 45: 337–344.

Medina, Ernesto. 1999. Mangrove physiology: the challenge of salt, heat, and light stress under recurrent flooding. Ecosistemas de manglar en América tropical. Mexico: Instituto de Ecología, AC, Costa Rica: UICN/ORMA, Silver Spring MD: NOAA/NMFS: 109–126.

Medina, Ernesto, Elvira Cuevas, and Ariel E Lugo. 2010. Nutrient relations of dwarf Rhizophora mangle L. mangroves on peat in eastern Puerto Rico. Plant ecology 207. Springer: 13–24.

Moraes, R M, W B C Delitti, and JAPV Moraes. 2003. Gas exchange, growth, and chemical parameters in a native Atlantic forest tree species in polluted areas of Cubatao, Brazil. Ecotoxicology and Environmental Safety 54. Elsevier: 339–345.

De Nicola, F, G Maisto, M V Prati, and A Alfani. 2008. Leaf accumulation of trace elements and polycyclic aromatic hydrocarbons (PAHs) in Quercus ilex L. Environmental Pollution 153. Elsevier: 376–383.

Office for Coastal Management. 2017. C-CAP Land Cover, Puerto Rico, 2010. Charleston, SC.

Paerl, H W, K L Webb, J Baker, and W J Wiebe. 1981. Nitrogen fixation in waters. Nitrogen fixation 1: 193–240.

Palma, Estibaliz, Jane A Catford, Richard T Corlett, Richard P Duncan, Amy K Hahs, Michael A McCarthy, Mark J McDonnell, Ken Thompson, Nicholas S G Williams, and Peter A Vesk. 2017. Functional trait changes in the floras of 11 cities across the globe in response to urbanization. Ecography 40. Wiley Online Library: 875–886.

Penuelas, Josep, and Iolanda Filella. 2002. Metal pollution in Spanish terrestrial ecosystems during the twentieth century. Chemosphere 46. Elsevier: 501–505.

Qiu, Yao-Wen, Ke-Fu Yu, Gan Zhang, and Wen-Xiong Wang. 2011. Accumulation and partitioning of seven trace metals in mangroves and sediment cores from three estuarine wetlands of Hainan Island, China. Journal of Hazardous Materials 190. Elsevier: 631–638.

Raciti, Steve Michael, P M Groffman, and T J Fahey. 2008. Nitrogen retention in urban lawns and forests. Ecological Applications 18. Wiley Online Library: 1615–1626.

Reich, P B, D S Ellsworth, and M B Walters. 1998. Leaf structure (specific leaf area) modulates photosynthesis–nitrogen relations: evidence from within and across species and functional groups. Functional Ecology 12. Wiley Online Library: 948–958.

Reich, Peter B, David S Ellsworth, Michael B Walters, James M Vose, Charles Gresham, John C Volin, and William D Bowman. 1999. Generality of leaf trait relationships: a test across six biomes. Ecology 80. Wiley Online Library: 1955–1969.

Reinking, Larry. 2007. Examples of Image Analysis Using ImageJ.

Reis, Carla Roberta Gonçalves, Gabriela Bielefeld Nardoto, and Rafael Silva Oliveira. 2017. Global overview on nitrogen dynamics in mangroves and consequences of increasing nitrogen availability for these systems. Plant and soil 410. Springer: 1–19.

Reis, Carla Roberta Gonçalves, Gabriela Bielefeld Nardoto, André Luis Casarin Rochelle, Simone Aparecida Vieira, and Rafael Silva Oliveira. 2017. Nitrogen dynamics in subtropical fringe and basin mangrove forests inferred from stable isotopes. Oecologia 183. Springer: 841–848.

Romero, Isabel C, Myrna Jacobson, Jed A Fuhrman, Marilyn Fogel, and Douglas G Capone. 2012. Long‐term nitrogen and phosphorus fertilization effects on N2 fixation rates and nifH gene community patterns in mangrove sediments. Marine ecology 33. Wiley Online Library: 117–127.

Savage, Candida. 2005. Tracing the influence of sewage nitrogen in a coastal ecosystem using stable nitrogen isotopes. AMBIO: A Journal of the Human Environment 34. BioOne: 145–150.

Singh, R P, and M Agrawal. 2007. Effects of sewage sludge amendment on heavy metal accumulation and consequent responses of Beta vulgaris plants. Chemosphere 67. Elsevier: 2229–2240.

Da Souza, Iara, Marina Marques Bonomo, Mariana Morozesk, Lívia Dorsch Rocha, Ian Drumond Duarte, Larissa Maria Furlan, Hiulana Pereira Arrivabene, Magdalena Victoria Monferrán, Silvia Tamie Matsumoto, and Camilla Rozindo Dias Milanez. 2014. Adaptive plasticity of Laguncularia racemosa in response to different environmental conditions: integrating chemical and biological data by chemometrics. Ecotoxicology 23. Springer: 335–348.

Suess, Hans E. 1955. Radiocarbon concentration in modern wood. Science 122. JSTOR: 415–417.

Takemura, Taro, Nobutaka Hanagata, Koichi Sugihara, Shigeyuki Baba, Isao Karube, and Zvy Dubinsky. 2000. Physiological and biochemical responses to salt stress in the mangrove, Bruguiera gymnorrhiza. Aquatic Botany 68. Elsevier: 15–28.

Tomašević, M, Z Vukmirović, S Rajšić, M Tasić, and B Stevanović. 2005. Characterization of trace metal particles deposited on some deciduous tree leaves in an urban area. Chemosphere 61. Elsevier: 753–760.

U.S. Census Bureau. 2015. 2015 TIGER/Line Shapefiles roads.

United States Army Corps of Engineers. 2015. Final Environmental Impact Statement Cano Martín Pena Ecosystem Restoration Project San Juan, Puerto Rico.

United States Census Bureau. United States Census Bureau Decennial Census of Population and Housing.

United States Census Bureau. 1913. Census of Population and Housing 1910.

United States Environmental Protection Agency. 1996. Method 3050B Acid digestion of sediments sludges, and soils revision 2.

Van der Valk, A G, and P M Attiwill. 1984. Acetylene reduction in an Avicennia marina community in Southern Australia. Australian journal of botany 32. CSIRO: 157–164.

Verma, Amitosh, and S N Singh. 2006. Biochemical and ultrastructural changes in plant foliage exposed to auto-pollution. Environmental Monitoring and Assessment 120. Springer: 585–602.

Webb, Richard M T, and Fernando Gómez-Gómez. 1998. Synoptic survey of water quality and bottom sediments, San Juan Bay estuary system, Puerto Rico, December 1994-July 1995. US Department of the Interior, US Geological Survey.

Weis, Judith S, and Peddrick Weis. 2004. Metal uptake, transport and release by wetland plants: implications for phytoremediation and restoration. Environment international 30. Elsevier: 685–700.

Westoby, Mark, and Ian J Wright. 2006. Land-plant ecology on the basis of functional traits. Trends in ecology & evolution 21. Elsevier: 261–268.

Wickham, Hadley. 2009. ggplot2: Elegant Graphics for Data Analysis. New York, NY: Springer-Verlag.

Williams, Nicholas S G, Mark W Schwartz, Peter A Vesk, Michael A McCarthy, Amy K Hahs, Steven E Clemants, Richard T Corlett, Richard P Duncan, Briony A Norton, and Ken Thompson. 2009. A conceptual framework for predicting the effects of urban environments on floras. Journal of ecology 97. Wiley Online Library: 4–9.

Wilson, J Bastow. 1988. A Review of Evidence on the Control of Shoot: Root Ratio, in Relation to Models. Annals of Botany 61: 433–449.

Yan, Jun, Emiliano A Valdez, Pravin K Trivedi, David M Zimmer, Andy Staudt, Arkady E. Shemyakin, Heekyung Youn, et al. 2011. R: A Language and Environment for Statistical Computing. R Foundation for Statistical Computing. Vol. 1. Vienna, Austria: R Foundation for Statistical Computing. doi:10.1007/978-3-540-74686-7.

Ye, Yong, Nora F Y Tam, Y S Wong, and C Y Lu. 2003. Growth and physiological responses of two mangrove species (Bruguiera gymnorrhiza and Kandelia candel) to waterlogging. Environmental and Experimental Botany 49. Elsevier: 209–221.

Zuberer, D_A, and W S Silver. 1978. Biological dinitrogen fixation (acetylene reduction) associated with Florida mangroves. Applied and environmental Microbiology 35. Am Soc Microbiol: 567–575.

